# Antigen-dependent activation of marginal zone B cells amplifies hypertension

**DOI:** 10.1101/2025.10.17.682980

**Authors:** Hericka B. Figueiredo Galvao, Maria Jelinic, Maggie Lieu, Vivian Tran, Buddhila Wickramasinghe, Jake N. Robertson, Tayla A. Gibson Hughes, Henry Diep, Asha Haslem, Mathew G. Lewsey, Alexander Bobik, Christopher G. Sobey, Tomasz J. Guzik, Grant R. Drummond, Antony Vinh

## Abstract

**Aims:** B cells contribute to the development of hypertension, yet, the specific B cell subsets involved, the mechanism underlying their activation, and the relevance of these responses to human disease remain poorly defined.

**Methods and results:** We used single-cell RNA sequencing, single-cell B cell receptor (BCR) VDJ sequencing, and high dimensional flow cytometry to characterise B cell responses in murine angiotensin II-induced hypertension. Chronic angiotensin II infusion in male and female mice increased systolic blood pressure and selectively expanded marginal zone B (MZB) cells, with evidence of antigen-dependent activation, including clonal BCR expansion, enrichment of IGHV1 B cell receptor variants, and increased expression of activation markers (CD69 and Nur77). Intercellular communication analyses revealed enhanced antigen-presentation signalling between MZB and CD8+ T cells in hypertensive mice. Activated MZB-like memory B cells also accumulated in the kidneys of hypertensive mice. Consistent with these findings, multiomic analysis of kidneys from patients with hypertensive chronic kidney disease (CKD) demonstrated an increase in memory B cells with a MZB phenotype and enrichment of antigen-presentation-linked communication with CD8+ T cells. Importantly, hypertensive responses to angiotensin II infusion were significantly blunted in mice lacking MZB cells (BAFF-R^-/-^).

**Conclusion:** Our findings identify MZB cells as a selectively activated, antigen-responsive B cell subset that amplified pathogenic immune responses in murine and human hypertension. By linking subset-specific BCR activation to immune cross-talk and disease causality, this study identifies MZB cells – and the (auto)antigens that activate them – as promising targets for precision immunomodulatory strategies in hypertension.

## 1. Introduction

The immune system plays a central role in the development of cardiovascular diseases including hypertension^1, 2^. Clinical studies from over 50 years ago linked hypertension with B cell activation, evidenced by elevated circulating IgG antibodies^3–6^. More recently, preclinical studies in experimental models of hypertension provide causal support for these associations, whereby genetic or pharmacological depletion of B cells reduces blood pressure elevation. Importantly, B cell depletion also limits cardiac and vascular remodelling, identifying B cells as key contributors to the development of hypertension and its sequalae^7–9^.

The findings above imply that B cells could be therapeutically targeted to treat hypertension. However, global B cell depletion is unlikely to be clinically viable due to its detrimental effects on immune function and infection control. More selective strategies, such as inhibiting the specific B cell subsets activated in hypertension or targeting the mechanisms that drive their activation may offer greater translational potential. In contrast to the extensive work defining the roles and pathogenic subsets of other immune cells in hypertension^1, 2^, including T cells and macrophages, relatively little effort has been devoted to achieving subset-level and mechanistic resolution for B cells. Thus, it remains unclear which B cell subsets drive hypertension, whether their activation is antigen-dependent, and how this knowledge might be therapeutically exploited.

Mature B cells originate from one of two lineages. The B-1 lineage, which includes B-1a and B-1b B cells, is the first to develop, appearing during foetal development^10^. B-1a and B-1b cells respond to T cell-independent antigens through one of two main mechanisms: (1) activation of innate cell surface immune receptors such as toll-like receptors, in type 1 T cell-independent responses^10^; or (2) recognition of phylogenetically conserved pathogenic sequences by B cell receptors (BCRs) in type 2 T cell-independent responses^10^. The B-2 lineage, which includes follicular (FO) and marginal zone (MZB) B cells, develops postnatally following the hematopoietic switch from the foetal liver to the bone marrow^10, 11^. FO B cells are primarily activated by antigens binding to their BCRs and require co-stimulation from T cells to exert their full response^11^. In contrast, MZB cells occupy a specialized niche at the interface of innate and adaptive immunity and display broader functional plasticity, responding efficiently to both T cell–independent and T cell–dependent antigens^11^.

With the exception of B-1a cells^10^, productive B cell activation leads to the expansion of the activated population. When this activation is antigen-specific, the resulting B cells express identical BCRs, also known as B cell clones. This process reduces overall BCR diversity and skews the repertoire toward specific immunoglobulin heavy chain variable regions (IGHVs)^12^.

In this study, we combined single-cell transcriptomics, BCR sequencing and spectral flow cytometry to: (1) identify which B cell subsets are activated during the development of angiotensin II-dependent hypertension in mice; (2) investigate the mechanisms underlying their activation; and (3) define the functional consequences of this activation. We observed that MZB cells were selectively expanded in the spleens of hypertensive mice and that a higher proportion of MZB cells expressed markers of activation. This expansion was accompanied by clonal BCR enrichment evidenced by an over-representation of IGHV1 BCR variants, consistent with antigen-dependent activation. Activated MZB-like cells also accumulated in the kidneys of hypertensive mice and in humans with hypertensive chronic kidney disease, where they exhibited a memory phenotype. Finally, mice deficient in MZB cells (BAFF-R^-/-^mice) were protected from hypertension, supporting a causal role for MZB cells. Together, these findings identify MZB cells as key mediators of hypertension, highlight their cross-species relevance, and provide a foundation for future studies aimed at identifying the antigens responsible for MZB cell activation.

## 2. Methods

Comprehensive methods and relevant references are included in the **Supplementary material, *Expanded Materials and Methods***.

### 2.1. Animals and ethics

All procedures were approved by the La Trobe University Animal Ethics Committee (AEC 16-93), the Monash University Animal Research Platform Ethics Committee (MARP/2016/077) and the Institutional Biosafety Committee (GM16-25). All experiments were performed in accordance with the Australian Code for the Care and Use of Animals for Scientific Purposes (8th Edition 2013, updated 2021).

Male and female C57BL6 mice aged 8- to 11-weeks were sourced from the La Trobe Animal Research and Teaching Facility (LARTF; Bundoora, Australia) and Monash Animal Services (Clayton, Australia). B cell activating factor-receptor knockout (BAFF-R^-/-^) mice were bred and housed in the Animal Research laboratories (Clayton, Australia) and LARTF (Bundoora, Australia). Mice were housed in temperature (22 ± 2°C) and humidity (55 ± 15%) regulated rooms with 12-hour light/dark cycles in individually ventilated cages (IVC) with access to food and water *ad libitum*. Hypertension was induced via subcutaneous infusion of angiotensin II (0.7 mg/kg/day) using osmotic minipumps (Alzet Model 2004, USA). Control mice received vehicle (0.1% acetic acid in saline). Infusions were maintained for either 14 or 28 days, as previously described^13^. Systolic blood pressure (SBP) was measured using tail-cuff plethysmography (MC4000 Multichannel System; Hatteras Instruments, USA), as previously described^14^. Mice were euthanised by carbon dioxide asphyxiation, then bone marrow, spleen, blood and kidney samples were harvested for preparation into single-cell solutions for high-dimensional (spectral) flow cytometry and/or single cell multiomic (RNA+VDJ) sequencing.

### 2.2. Single-cell and single-nuclear RNA sequencing of human kidneys

Multiomic sequencing data of healthy, diabetic and hypertensive human kidneys were downloaded from GSE211785^15^. Samples from control and hypertensive chronic kidney disease (HKD) participants were then extracted for downstream analyses.

### 2.3. Statistical analyses

Power calculations determined that a minimum sample size of n = 9 was required to detect a 25% change in mean SBP with a standard deviation of 10%, 80% power and α = 5%.

All physiological data analyses were performed using GraphPad Prism 9.4.0 (GraphPad Software, San Diego, California, USA). Data distribution was assessed using the D’Agostino & Pearson test, and outliers were identified using a ROUT method (false discovery rate = 1%). Data are presented as the mean ± standard error of the mean (SEM). Results were considered statistically significant when *p* < 0.05. SBP data were analysed using a two-way mixed effects model with Geisser-Greenhouse correction and Tukey’s multiple comparisons test. Flow cytometry data was analysed using a two-tailed unpaired t-test. For sex- or genotype-specific analyses, SBP datasets were analysed using a three-way mixed effects model with Geisser-Greenhouse correction, followed by Tukey’s *post hoc* test. All other physiological data were analysed using a two-way ANOVA followed by Tukey’s *post hoc* test.

For bioinformatics analyses, differentially expressed genes (log_2_(fold change) ≥ 0.25 and expressed by ≥ 10% of cells) were analysed using Wilcoxon rank-sum tests comparing each cluster to all others. Genes with a Bonferroni-adjusted *p* ≤ 0.01 were considered significant. For cell type predictions via SingleR, differentially expressed genes were evaluated using a Wilcoxon rank-sum test (with adjusted *p* < 0.05). For gene ontology (GO) enrichment significance was assessed using the Benjamini-Hochberg correction (*q* < 0.05).

## 3. Results

### 3.1. Single-cell sequencing of primary and secondary lymphoid organs identifies several lymphoid and non-lymphoid cell populations

Single-cell suspensions of bone marrow and spleen samples from male and female mice infused with either angiotensin II or vehicle were subjected to transcriptomic sequencing. Unsupervised clustering identified a total of 8 non-lymphoid and 3 lymphoid populations at the broad lineage level: pro-neutrophils, neutrophils, erythroid primed monocyte-erythrocyte progenitors (E-MEPs), granulocyte-monocyte progenitors (GMPs), pro-monocytes, monocytes, basophils, dendritic cells (DCs), natural killer cells (NKs), T cells and B cells (**Fig. 1A**). The key genes used to identify each of these populations and their top 5 differentially expressed genes are shown in **Fig. 1B** and **C**, respectively. Relevant references for the main population gene markers are available in ***Table S5***.

**Figure 1:**
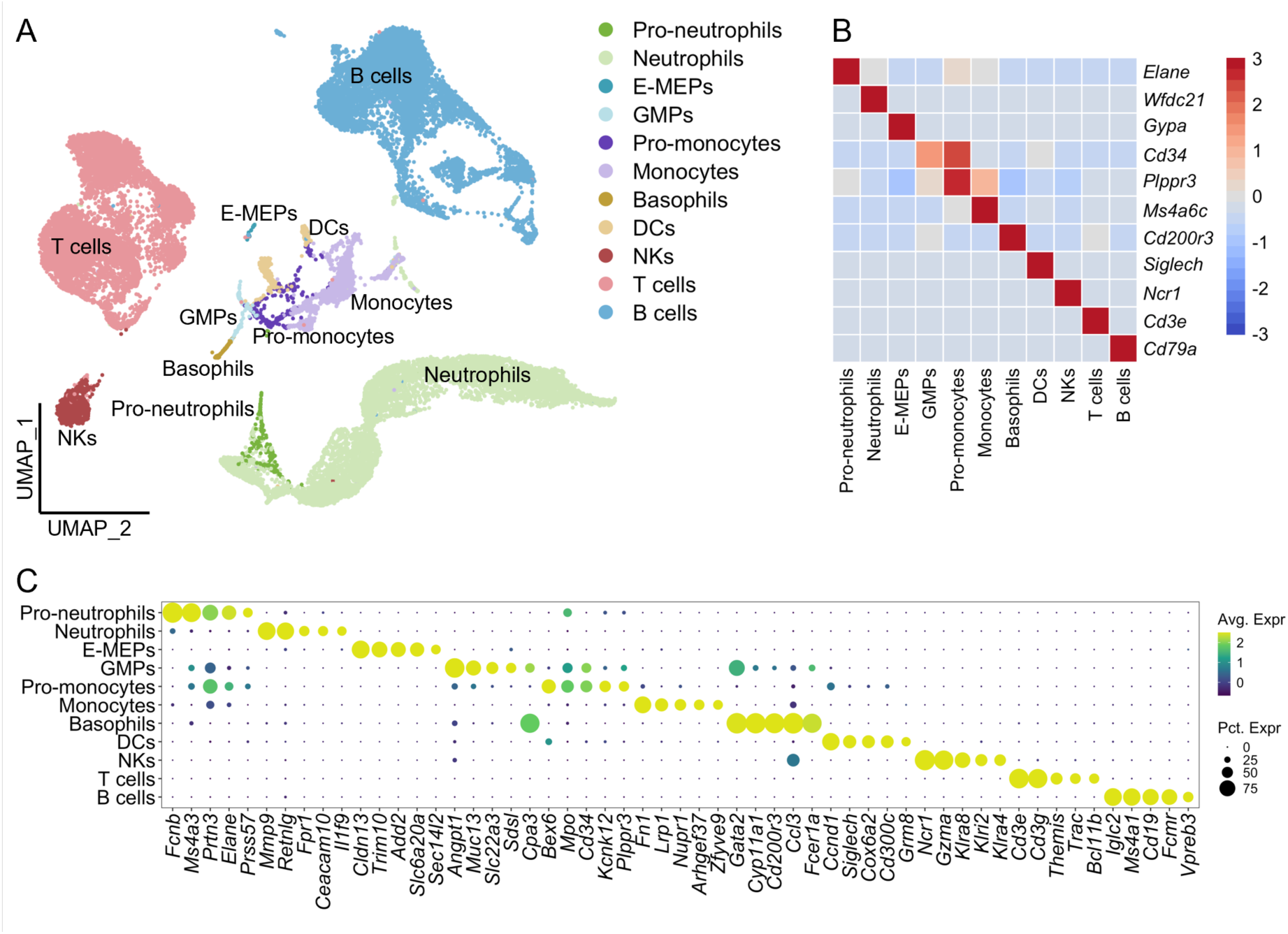
Hematopoietic cell types identified by single cell transcriptomics in the bone marrow and spleen. A) UMAP dimensional reduction of all cells identified in the bone marrow and spleen. B) Heatmap showing the expression of key cell markers from low (blue) to high (red) expression. C) Top 5 uniquely expressed genes for each population where the size and colour of each dot represents the proportion of cells expressing a given gene in that cluster and its normalized expression count, respectively. E-MEPs= erythroid primed monocyte-erythrocyte progenitors; GMPs = granulocyte-monocyte progenitors; DCs = dendritic cells; NKs = natural killer cells

### 3.2. Gene ontology analysis of spatially correlated genes corroborates B cell subpopulation classification

B cells were further classified into subpopulations, each defined by at least 3 differentially expressed marker genes. In total, 15 B cell populations were identified across bone marrow and spleen samples (**Fig. 2A-B**). To corroborate B cell subtype identification, a three-dimensional pseudotime trajectory was generated (**Fig. 2C**) and the genes enriched along the trajectory were grouped into gene modules (**Fig. 2D**) for GO analyses (**Fig. 2E-I**). Relevant references for B cell subpopulation gene markers are available in ***Table S6***.

**Figure 2:**
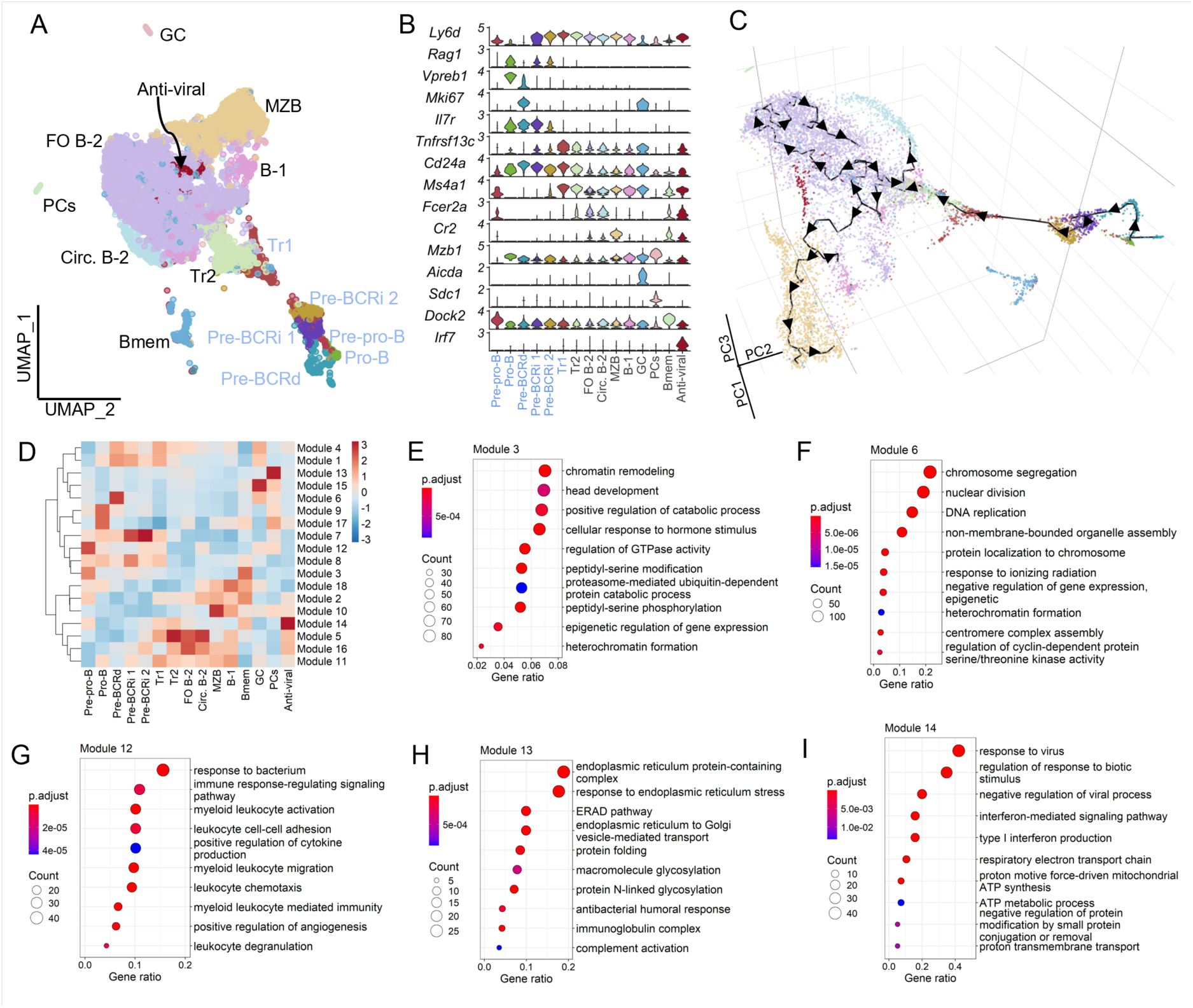
Pseudotime trajectory projection and gene ontology modules corroborate with identified B cell subtypes. A) UMAP dimensional reduction of B cell sub-clustering. B) Violin plots of genes used to identify B cell developmental stages and subtypes. C) Three-dimensional (3D) pseudotime trajectory (black trace with arrows) showing the maturation of B cells from the bone marrow to the spleen. D) Genes expressed in specific regions of the 3D space were grouped into modules and visualised using a heatmap. Dot plots of gene ontologies from genes expressed in module 3 (E), module 6 (F), module 12 (F), module 13 (H) and module 14 (I) were obtained to identify cell functions. In panels E-I, the size and colour of each dot represents the number of genes expressed in a given ontology and its adjusted P value, respectively. Immature B cell populations found exclusively in the bone marrow are shown in cornflower blue. Pre-BCRd = pre-B cell receptor dependent cells; Pre-BCRi = pre-B cell receptor independent cells; Tr1 = transitional B cells found in bone marrow; Tr2 = transitional B cells found in spleen; MZB = marginal zone B cells; Bmem = memory B cells; GC = germinal center B cells; PCs = plasma cells; Circ. B-2 = circulating B-2 cells.

Low expression of *Ly6d* in the pre-pro B, pro-B and pre-B cell receptor dependent (pre-BCRd) populations indicates their recent commitment to a B cell fate (**Fig. 2B; *Table S6***). Pro-B cells were identified by their additional expression of *Rag1*, *Rag2* and the BCR surrogate light chains *Vpreb1*, *Vpreb2* and *Vpreb3* (**Fig. 2B; *Table S6; Fig. S4A***). Large pre-B cells (pre-BCRd) receive tonic signals from the pre-BCR to proliferate and were characterised by the expression of immature BCR surrogate light chains and proliferation-associated genes (*Mki67*, *Pclaf* and *Top2a*; **Fig. 2B; *Table S6; Fig. S4A, B***). Consistent with their proliferative phenotype and active cell cycling, Pre-BCRd cells were enriched with genes from Module 6 which contained GO terms related to proliferation (*DNA replication*, *nuclear division* and *chromosome segregation*; **Fig. 2D, F**).

As shown previously^16^, we identified two small pre-B cell populations: pre-BCR-independent- 1 (pre-BCRi 1) and pre-BCR-independent-2 (pre-BCRi 2). Both populations lacked expression of proliferative genes but expressed low levels of mature BCR light chain genes (*Iglc1*, *Iglc2* and *Iglc3*), indicating ongoing light chain rearrangement (**Fig. 2B; *Table S6; Fig. S4B***). Consistent with their immature phenotypes, the IL7 receptor (*Il7r*) was enriched in the pro-B, pre-BCRd and pre-BCRi 1 subpopulations (**Fig. 2B; *Table S6***). *IL7r* expression was absent in mature B cells and lower in the pre-BCRi 2 population, suggesting a more differentiated state in the latter compared to the pre-BCRi 1 population (**Fig. 2B**).

Transitional B cells in the bone marrow, annotated here as Tr1, expressed the B cell activating factor receptor (BAFF-R; *Tnfrsf13c*), which provides them with survival signals, as well as high levels of *Cd24a*, a common marker of transitional B cells (**Fig. 2B; *Table S6***). Transitional B cells in the spleen (Tr2) closely resembled Tr1 cells in the bone marrow, as evidenced by their shared expression of transcriptional regulators (**Fig. 2B; *Table S6; Fig. S4A-C***) and differentially expressed genes (***Table S6;* Fig. S4D**).

Beyond the transitional stages, mature B cells with distinct phenotypic and functional properties were also identified in our dataset. These included follicular B-2 (FO B-2), marginal zone B cells (MZB), circulating B-2 (Circ. B-2), B-1 B cells, germinal center (GC), memory (Bmem) and plasma cell (PC) populations were also identified in our dataset.

FO B-2 cells in the spleen were defined by the expression of *Fcer2a*, *Cr2* (CD21) and the transcription factor *Klf2*, known to maintain the follicular phenotype (**Fig. 2B; *Table S6; Fig. S4C-D***). MZB cells were distinguished from FO B-2 cells, by the high expression of *Cr2* and *Mzb1*, with low levels of *Klf2* and *Fcer2a* (**Fig. 2B; *Table S6; Fig. S4C-D*)**. Following maturation in the spleen, B cells may re-enter the circulation as mature circulating B cells. In the bone marrow, we identified a B cell population lacking the expression of immature markers and expressing mature B-2 lineage markers (*Fcer2a* and *Cr2;* **Fig. 2B; *Table S6; Fig. S4A***) that we describe here as circulating B-2 cells. B-1 cells were identified by enriched *Mzb1* expression and the absence of B-2 lineage markers (*Fcer2a* and *Cr2*), consistent with their distinct developmental origin and functional profile (**Fig. 2B; *Table S6; Fig. S4C***).

During germinal center (GC) reactions, activated FO B-2 and MZB cells receive co-stimulatory signals from T cells and differentiate into either memory B cells or plasma cells. We identified a small population of GC cells expressing proliferation markers (*Mki67*, *Pclaf* and *Top2a*) and activation-induced cytidine deaminase (*Aicda*), which initiates somatic hypermutation and class-switch recombination of BCRs (**Fig. 2B; *Table S6; Fig. S4B***). In addition to Module 6 (proliferation/expansion GO terms), GC cells were also enriched for Module 12 (**Fig. 2D,G**), which is associated with *leukocyte cell-cell adhesion*, *immune response-regulating signalling pathway*, *and positive regulation of cytokine production*, supporting their role in receiving co-stimulatory signals from T cells.

A population of mature B cells expressing high levels of *Bach2* and *Dock2,* required for memory B cell differentiation, was identified and labelled as memory B cells (**Fig. 2B; *Table S6; Fig. S4D***). Consistent with a memory phenotype, this population was enriched for GO terms associated with *epigenetic regulation of gene expression*, *chromatin remodelling* and the *maintenance of stem cell populations* (Module 3; **Fig. 2D,E**).

Plasma cells (PCs) were also identified in the spleen and bone marrow by their expression of the plasma cell-specific B cell maturation antigen receptor (BCMA; *Tnfrsf17*), the transcription factor B lymphocyte-induced maturation protein 1 (Blimp-1; *Prdm1*) and *Sdc1* (**Fig 2B; *Table S6; Fig. S4C*).** PCs are professional antibody-secreting cells and accordingly their GO terms were largely associated with *secretory pathways*, *protein folding* and *humoral immune responses* (Module 13; **Fig. 2D,H**). An additional distinct population of mature B cells expressing interferon-stimulated genes, including *Ifit3, Isg15, Irf7* (**Fig. 2B; *Table S6; Fig. S4C***), with gene ontologies associated with *anti-viral responses* and *interferon-mediated signalling* (Module 14; **Fig. 2D,I**), was tentatively annotated as “anti-viral” B cells.

### 3.3. Expansion and activation of marginal zone B cells in spleens of chronically hypertensive mice

Angiotensin II infusion increased systolic blood pressure in mice from a baseline of 119.8 ± 3.6 mmHg to 167.2 ± 2.6 mmHg by day 28 (**Fig. 3A**). In contrast, there was no change in blood pressure across the 28 days in vehicle-infused control mice. Comparative analysis of transcriptomic data from normotensive and hypertensive mice revealed a selective expansion of marginal zone B (MZB) cells, with no other B-cell subset showing a comparable increase. The proportion of MZB cells was higher in hypertensive cf. normotensive mice (21.5 ± 2.5% vs 16.8 ± 0.9% of sequenced splenic B cells, respectively; **Fig. 3B, C**). This selective expansion was independently confirmed by flow cytometry, which demonstrated a higher frequency of MZB cells among live splenocytes in hypertensive mice (5.4 ± 0.3% vs 4.2 ± 0.2%; **Fig. 3D**). In addition to increased abundance, splenic MZB cells from hypertensive mice exhibited elevated expression of the activation marker CD69 (**Fig. 3E**), indicating that this population is not only expanded but also activated in hypertension.

**Figure 3:**
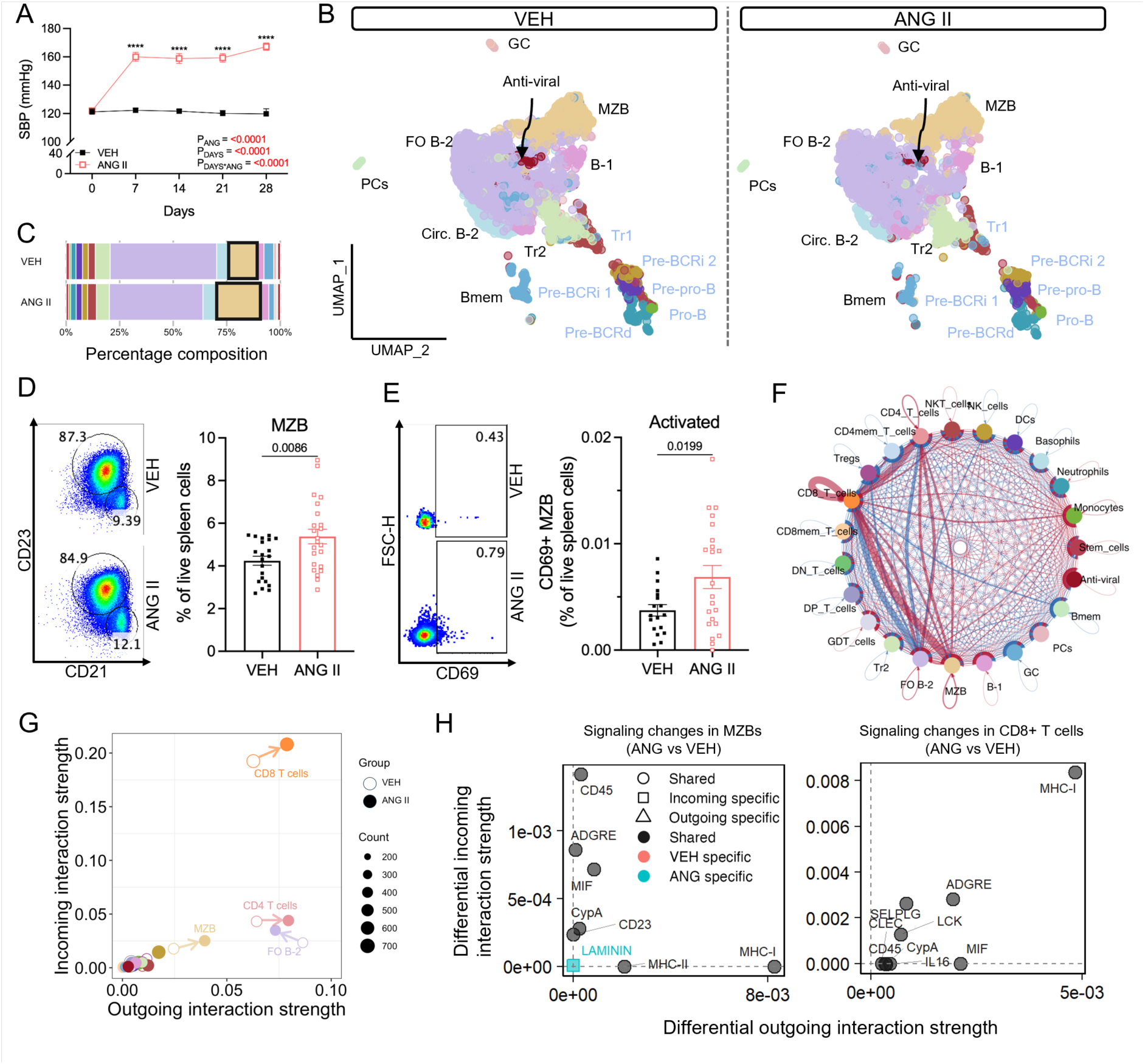
Expansion and activation of marginal zone B cells in hypertensive mice. A) Systolic blood pressures in vehicle (0.5% NaCl, 0.1% acetic acid) and angiotensin II (0.70 mg/kg/day)-infused mice. B) UMAP dimensional reduction of B cells from normotensive and hypertensive mice split by treatment. C) Bar graph showing the relative proportion of B cells as percentages in the two conditions. Marginal zone B cells are highlighted by a black box. Bar graph and corresponding flow cytometry plots showing total (D) and activated (E) CD21^hi^CD23^neg^ MZB, as a percentage of all live spleen cells in both conditions. F) Changes in intercellular communication networks in response to angiotensin II-infusion, where the line thickness indicates the relative interaction strength and the colour its direction relative to vehicle-infused mice (red = increased; blue = decreased). G) Scatter plot of incoming and outgoing interactions for each cell population. Vehicle- and angiotensin II-infused mice are shown by open and closed circles, respectively. The size of each circle represents the total number of incoming and outgoing interactions. Arrows highlight an increase or decrease in interactions from the same cluster between the two conditions. H) Enriched signalling pathways in MZB and CD8+ T cells from hypertensive mice relative to their normotensive counterparts. X-axes show increases in outgoing signalling, and y-axes increases in incoming signalling. For A), D) and E) data are represented as the mean ± standard error of the mean. Systolic blood pressures were analysed using a two-way ANOVA. **** indicates a P value <0.0001. Treatment effects are indicated by P_ANG_ for angiotensin II, P_DAYS_ for time, and P_DAYS*ANG_ for interaction effects. Flow cytometry data was analysed using a two-tailed unpaired student’s T test. P values are indicated above bar graphs. Immature B cell populations found exclusively in the bone marrow are shown in cornflower blue. Pre-BCRd = pre-B cell receptor dependent cells; Pre-BCRi = pre-B cell receptor independent cells; Tr1 = transitional B cells found in bone marrow; Tr2 = transitional B cells found in spleen; MZB = marginal zone B cells; Bmem = memory B cells; GC = germinal center B cells; PCs = plasma cells; Circ. B-2 = circulating B-2 cells.

To determine how the selective expansion of MZB cells alters immune cross-talk in hypertension, we compared ligand-receptor expression across splenic cell clusters from normotensive and hypertensive mice. We observed a marked increase in communication strength between MZB cells and CD8^+^ T cells in hypertensive mice (**Fig. 3F**). Consistent with increased communication strength, a comparison of the number of inferred communication links and their communication probability revealed an increase in both incoming and outgoing communication links for MZBs and CD8+ T cells (**Fig. 3G**). We next applied manifold and classification learning to identify specific signalling pathways that were differentially expressed between the two conditions. Outgoing antigen presenting pathways (MHC-I and MHC-II) were increased in MZB cells from hypertensive mice (**Fig. 3H**). Interestingly, CD8^+^ T cells from hypertensive mice showed an increase in both outgoing and incoming antigen presentation (MHC-I; **Fig. 3H**). Collectively, these analyses suggest an increase in antigen-presentation between these two immune cell populations.

Given the well-recognised sexual dimorphism in angiotensin II-dependent hypertension, we examined whether MZB cell expansion and activation differed between male and female mice. Although angiotensin II infusion increased SBPs in both sexes, the pressor response in female mice was delayed and, in contrast to males, was not accompanied by cardiac hypertrophy (***Fig. S5***). Despite these physiological differences, MZB cell responses were remarkably similar across sexes. Both male and female mice exhibited comparable magnitudes of MZB cell expansion and activation (***Supplementary File 1***). In contrast, sex-specific differences were observed in other B cell subsets (***Supplementary File 1*).** Together, these findings suggest MZB cell activation and expansion in hypertension occurs independently of sex.

### 3.4. Marginal zone-like B cells accumulate in the kidneys and adopt a memory phenotype during chronic hypertension in mice

MZB cells typically reside in the marginal sinus of the spleen. However, MZB-like populations have been reported in inflamed tissues in autoimmune conditions such as thyroiditis^17^, diabetes^18^ and systemic lupus erythematosus^19^. In addition to the spleen and bone marrow, we extended our flow cytometry analysis to blood and kidney samples from hypertensive and normotensive mice. The kidneys are particularly relevant in hypertension, serving both as regulators of salt and water balance, and as a site susceptible to injury and dysfunction due to mechanical forces imposed by elevated pulse pressures^1^.

We found the proportion of total B cells in kidneys from hypertensive mice was almost double that in normotensive controls (2.5 ± 0.4% vs 1.4 ± 0.2% of live kidney cells; **Fig. 4A**). Most B cells in the kidneys exhibited a phenotype resembling that of MZB cells in the spleen, expressing CD19, B220 and CD21 with little to no expression of CD23 (**Fig. 4B**). Consistent with our findings in the spleen, these MZB-like cells showed increased expression of the activation marker CD69 in hypertensive mice (**Fig. 4C**). Notably, renal MZB-like cells in hypertensive mice exhibited a pronounced shift toward a memory phenotype, with approximately three-fold higher numbers of CD27⁺ cells compared with normotensive mice (**Fig. 4D**). A comprehensive summary of the B cell populations from all samples analysed by flow cytometry can be found in ***Supplementary File 2***.

**Figure 4:**
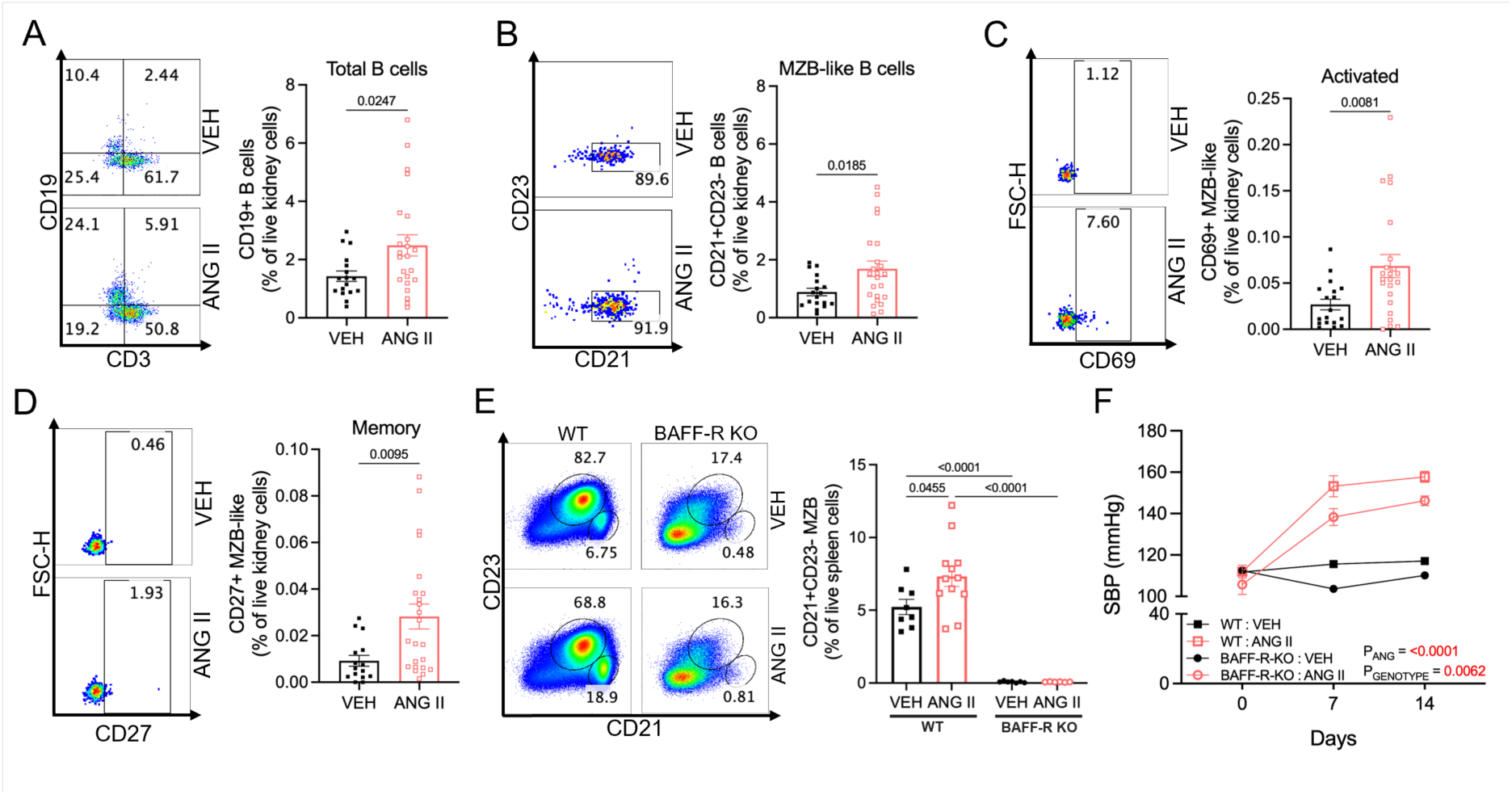
Marginal zone-like B cells infiltrate the kidneys of hypertensive mice, are activated and form memory B cells. Bar graphs and corresponding flow cytometry plots showing total B cells, MZB-like cells, activated MZB-like (CD69+), and memory MZB-like (CD27+) cells in panels A-D, respectively, in kidneys from normotensive and hypertensive mice. E) Bar graphs and corresponding flow cytometry plots of MZB cells in spleens of wildtype (WT) and B cell activating factor receptor knockout (BAFF-R KO) mice infused with either vehicle (0.5% NaCl, 0.1% acetic acid) or angiotensin II (0.70 mg/kg/day) solutions. B cells were identified as CD19+ and MZB-like cells as CD19^+^CD24^neg^B220^+^CD21^+^CD23^neg^ live cells in panels A-E. F) Systolic blood pressures of vehicle- and angiotensin II-infused WT and BAFF-R KO mice. All data are represented as the mean ± standard error of the mean. Flow cytometry data was analysed using a two-tailed unpaired student’s T test, with P values shown above bar graphs. Systolic blood pressures were analysed using a three-way ANOVA. Treatment effects are indicated by P_ANG_ for angiotensin II and P_GENOTYPE_ for genotypes.

### 3.5. Marginal zone B cell deficiency blunts angiotensin II-induced hypertension

To directly test the causal contribution of marginal zone B (MZB) cells in hypertension, we used BAFF-R^-/-^ mice, which lack mature B cells, including both MZB and FO B cells. BAFF-R^-/-^ and wild type mice were infused with angiotensin II, then B cell composition and blood pressure responses were assessed. In wild type mice, angiotensin II induced a robust increase in systolic blood pressure accompanied by expansion of MZBs in the spleen (**Fig. 4E, F**). In contrast, MZBs were virtually absent in spleens from BAFF-R^-/-^ mice under both basal (vehicle-infused) and hypertensive conditions (**Fig. 4E**) and these mice exhibited a markedly blunted pressor response to angiotensin II (**Fig. 4F**).

Although BAFF-R^-/-^ mice lack both MZB and FO B cells, FO B cells did not expand or exhibit evidence of activation in chronically hypertensive wild-type mice in any of the tissues examined by flow cytometry (***Supplementary File 1***). Furthermore, ligand-receptor analyses showed a reduction in FO B-2 communication strength (**Fig. 3F**) and outgoing communication probability (**Fig. 3G**), suggesting FO B-2 cells are less active in hypertensive mice cf. normotensive mice. Together, these findings indicate the attenuated hypertensive response observed in BAFF-R⁻/⁻ mice is attributable to the absence of MZB cells rather than FO B cells, supporting a direct pathophysiological role for MZB cells in hypertension.

### 3.6. Antigen-specific adaptations of B cell receptors of marginal zone B cells in hypertension

The increase in antigen-presenting signalling pathways (**Fig. 3H**) together with the accumulation of memory MZB cells in the kidneys (**Fig. 4D**), suggests that MZBs undergo antigen-specific activation in hypertension. Antigen-driven B cell activation is typically associated with reduced BCR diversity due to clonal expansion and preferential enrichment of specific heavy chains variants with a higher affinity for the activating antigen, a phenomenon known as BCR skewing^12^.

To assess this, we performed single-cell VDJ sequencing to quantify changes in BCR diversity, clonal expansion patterns and BCR heavy chain usage in MZBs. Both total BCR diversity and IgM diversity was reduced in MZBs from hypertensive mice, as assessed by Shannon entropy and Simpsons indices (**Fig. 5A, B**). Additionally, the number of expanded BCR clones in the spleens was increased in hypertensive mice (**Fig. 5C**). Hence, these changes are consistent with antigen-dependent activation.

**Figure 5:**
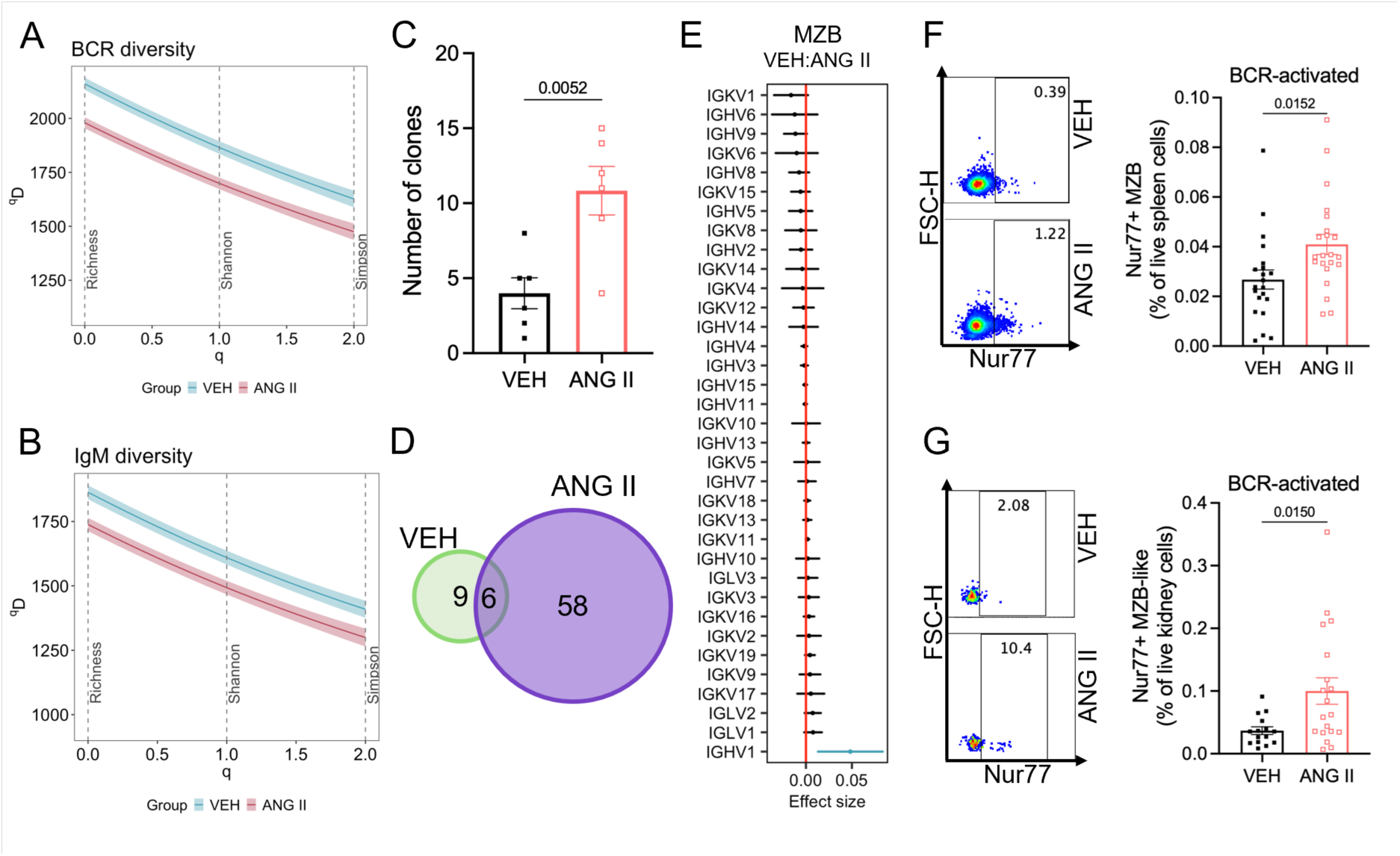
Reduced B cell receptor (BCR) diversity, BCR clonal expansion and increased expression of IGHV1 BCR variants in hypertension. Plots showing total BCR diversity (A) and IgM-only BCR diversity (B) in vehicle and angiotensin II-infused mice. Hill numbers (q) were used to calculate diversity (qD) metrics of BCR samples. Hill numbers can be interpreted over intervals where q = 0 represents species richness, q = 1 the Shannon Entropy of a population and q = 2 the Simpson Index, where Shannon and Simpson indices reflect species diversity. Within diversity curves, the darker solid line and its corresponding lighter band represent the mean and 95% confidence intervals, respectively. Lower qD values indicate reduced diversity and bands which do not overlap are statistically significant. C) Bar graph showing the number of expanded B cell clones in sequenced spleens. D) Venn diagram showing number of overlapping clones in normotensive and hypertensive mice. E) Mean effect size changes of the treatment effect (VEH:ANG II) on the frequency of variable gene segments in antibody heavy and light chains from the spleen. Bar graphs and corresponding flow cytometry plots showing expression of the BCR-specific activation marker Nur77 in MZBs in spleens (F) and kidneys (G) from normotensive and hypertensive mice. For C, F and G, data are represented as the mean ± standard error of the mean, where data was analysed using a two-tailed unpaired student’s T test. For E) hypothesis testing was performed using a two-way ANOVA. MZB = marginal zone B cells.

Although antigen specificity is encoded by all three complementarity-determining regions (CDR) of the BCR, the third CDR (CDR3) loop is the most diverse and represents the main determinant of antigen specificity^20^. Comparative analysis of CDR3 amino acid sequences revealed 58 unique BCR clones in hypertensive mice that were absent from normotensive controls, present as either expanded clones or unexpanded BCRs (**Fig. 5D**). To determine whether BCR heavy-chain usage was also skewed, we quantified IGHV family usage in individual mice. Hypertensive mice had significantly higher proportions of MZB cells expressing IGHV1 BCR variants (**Fig. 5E**). Together, these data provide convergent evidence that MZB cells undergo antigen-dependent activation in hypertension and implicate IGHV1-expressing variants as key mediators of this response.

Nur77 is an immediate-early gene that is rapidly upregulated following direct BCR activation in both mice and humans^21^, thus serving as a surrogate marker of BCR-specific signalling. Using flow cytometry, we observed increased expression of Nur77 in MZBs from both the spleen (**Fig. 5F**) and kidneys (**Fig. 5G**) of hypertensive mice. Collectively, these findings identify MZB cells as the principal B cell subset activated in hypertension and provide independent functional evidence that this is likely antigen-driven.

### 3.7. Memory B cells with a MZB phenotype accumulate in kidneys of humans with hypertensive chronic kidney disease

Hypertension is a major driver of chronic kidney disease^22^. To test whether the renal B-cell phenotype observed in hypertensive mice is also evident in humans, we analysed published single-cell and single-nuclear RNA sequencing data from kidneys of patients with hypertensive chronic kidney disease (HKD)^15^.

Consistent with our mouse findings, HKD kidneys showed a significant increase in memory B cells (**Fig. 6A, B**). Notably, these memory B cells displayed an MZB-like transcriptional profile with high expression of CD21 and little to no expression of CD23 (**Fig. 6C**). Ligand-receptor analyses of HKD kidneys further mirrored the murine spleen network, revealing increased communication strength between memory B cells and CD8^+^ T cells in HKD kidneys relative to controls (**Fig. 6D**). Although overall interaction strength was lower than in the spleen, memory B cells in HKD kidneys exhibited increased outgoing antigen-presentation signalling (MHC I and MHC II), while CD8^+^ T cells also showed increased incoming and outgoing MHC I signalling (**Fig. 6E**). Together, these human data support the translational relevance on an MZB-like, antigen-presentation linked B cell program in hypertensive disease.

**Figure 6:**
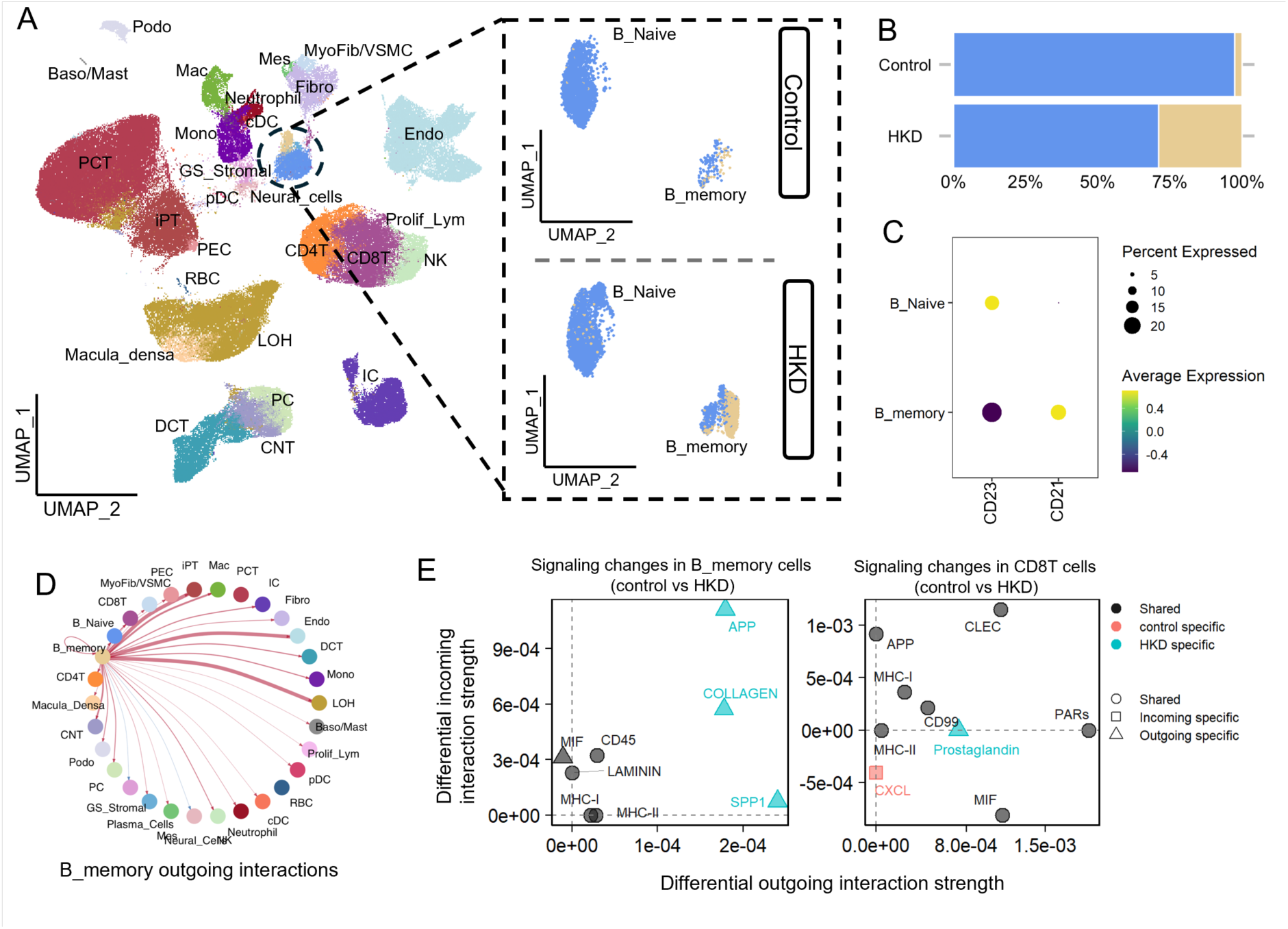
Memory B cells with a marginal zone B cell phenotype are expanded in samples from patients with hypertensive chronic kidney disease. A) UMAP dimensional reduction of cell populations identified in ^1515^, with an inset showing the increase in memory B cells in patients with hypertensive chronic kidney disease (HKD). B) Bar graph showing the relative proportion of B cells as percentages in the two conditions. C) Dot plot showing the expression of CD23 and CD21, where the size and colour of the dot represent the percentage of cells expressing each gene and its normalized average expression, respectively. D) Changes in outgoing memory B cell specific communication networks in patients with HKD, where the line thickness indicates the relative interaction strength and the colour its direction relative to control samples (red = increased; blue = decreased). E) Enriched signalling pathways in memory B cells and CD8+ T cells from HKD samples relative to their control counterparts. X-axes show increases in outgoing signalling, and y-axes increases in incoming signalling.

At a broader level, comparison of overall signalling changes identified 11 shared pathways that were upregulated in hypertensive mice and humans, including cell adhesion pathways (CADM, CDH1, Laminin, AGDRG, ICAM), TGF-β signalling, pro-inflammatory IL-16, and complement pathways (***Fig. S6A, B***).

## 4. Discussion

### Accumulating evidence supports a pro-hypertensive role for B cells^7–9^. However, the specific

B cell subsets involved and whether their activation is antigen-dependent have remained unclear. Here, using single-cell multiomic (RNA+VDJ) sequencing and high-dimensional flow cytometry, we identify marginal zone B (MZB) cells as the principal B-cell subset activated during angiotensin II-induced hypertension. MZBs exhibited clonal expansion and antigen/BCR-dependent activation in the spleen and kidney, forming a memory-like population in hypertensive kidneys. Importantly, parallel analyses of human hypertensive chronic kidney disease revealed an increase in memory B cells with an MZB phenotype and similarly implicated enhanced antigen-presentation signalling between MZB cells and CD8^+^ T cells. Finally, BAFF-R^-/-^ mice, which are deficient in MZB cells, displayed an attenuated hypertensive response to angiotensin II, supporting a direct pathophysiological role for this B cell subset in the development of hypertension.

MZB cells are a functionally versatile B cell population capable of responding to both T cell-dependent and T cell-independent antigens, enabling^11^ contributions to both innate and adaptive immune responses^11^. Although traditionally considered to be confined to the splenic marginal zone^23–25^, emerging evidence indicates that MZB cells can also localise to extra-splenic compartments, including the bone marrow, where they localise around sinusoidal capillaries and contribute to T cell-independent responses to blood-borne antigens^26^. In the present study, we identified MZB-like cells in the kidneys of hypertensive mice, characterised by expression of activation and memory markers. We further identified a comparable increase in memory B cells with an MZB phenotype in the kidneys of patients with hypertensive CKD. These findings indicate that MZB cells are not only activated in murine and human hypertension but also accumulate in renal tissues, positioning them to directly influence blood pressure regulation and disease pathogenesis^1^. Supporting this concept, previous studies have shown that genetic depletion of B cells reduces renal expression of the vasopressin 2 receptor, implicating B cells in mechanisms of salt and water handling^9^. Moreover, MZB cells have been identified within inflamed tissues in autoimmune diseases associated with hypertension, where they similarly exhibit activation and memory features^17–19^. Collectively, these observations support a role for MZB cells as tissue-resident immune modulators in hypertension and provide a rationale for further investigation of their contributions to renal dysfunction.

MZB cells possess several properties through which they may contribute to hypertensive disease progression. MZBs can modulate immune responses through cytokine secretion, thereby shaping the activity of other immune cell populations. For example, B cell-derived tumour necrosis factor-alpha promotes plaque formation and instability in atherosclerosis^27^, while interleukin-6 and interferon-gamma drive T cell proliferation and Th17 polarisation in autoimmune diseases^28, 29^. Notably, Th17 responses, characterised by the secretion of the pro-inflammatory cytokine, interleukin-17, have been implicated in both murine and human hypertension^1, 2^.

In addition to cytokine-mediated effects, MZB cells may contribute to vascular and renal dysfunction through antibody production. Elevated circulating and tissue levels of IgG antibodies are a consistent feature of clinical and experimental hypertension, and agonistic autoantibodies targeting the angiotensin II type 1^30–32^ and endothelin-1^33^ receptors have been reported in hypertension and pre-eclampsia. Although MZB cell activation is classically associated with the low-affinity IgM^+^ memory responses, these cells can undergo class switching to produce IgG and IgA^34^. In particular, IgG3 production is linked to T cell- independent activation by repetitive polysaccharide or complement-coated antigens^35, 36^, while IgA class switching can be driven by innate stimuli such as lipopolysaccharide (LPS) and TGF-β^37^. Consistent with these pathways, we have previously shown that angiotensin II-dependent hypertension in mice is associated with elevated levels of IgG3 antibodies in the circulation and vascular tissues^8, 38^ and here we observed *in silico* enrichment of TGF-β and complement signalling in both murine and human hypertension.

Beyond cytokine secretion and antibody production, MZB cells can act as antigen-presenting cells capable of amplifying adaptive immune responses. This function could be particularly relevant within hypertensive target organs such as the kidney, where sustained immune activation contributes to tissue injury and fibrosis. Following activation, MZB cells can migrate from the marginal zone into splenic follicles, where they present antigens to follicular dendritic cells and T cells^39, 40^. In the context of hypertension, we have previously shown that B cells exhibit increased expression of the co-stimulatory molecule CD86^8^ and that genetic or pharmacological disruption of B7-dependent co-stimulation attenuates T cell activation and increases in blood pressure^13^. Extending these findings, our ligand-receptor expression analyses indicate that MZBs engage in antigen-presentation signalling with CD8 T cells in both murine and human hypertension. This interpretation is further supported by multiomic profiling of human kidneys, which revealed enrichment of antigen-processing and presentation pathways and close spatial association of B and T cells with injured proximal tubule cells, within fibrotic microenvironments^15^. Collectively, these data support antigen presentation as a central mechanism through which MZB cells amplify pathogenic immune responses in hypertension.

Previous studies showed that RAG-1 knockout (RAG-1^-/-^) mice, which lack both B and T cells, are partially protected from experimental hypertension, highlighting a requirement for adaptive immunity in disease development^41^. However, adoptive transfer of B cells alone into RAG-1^-/-^ mice failed to restore the hypertensive phenotype, indicating that B cells require interaction with T cells to exert pro-hypertensive effects^41^. In contrast, genetic deletion of B cells alone resulted in a similarly attenuated hypertensive response that was reversible upon B cell reconstitution^8^. These observations are consistent with a model in which B cells do not act as independent effectors but instead amplify pathogenic T cell responses. In this context, MZB cells are well positioned to fulfil this role through antigen presentation. Interestingly, clinical hypertension is known to be associated with elevated circulating complement levels^42–44^ and complement activation has been implicated in multiple models of hypertension^45–48^ ^49^. Because MZB cells are highly responsive to complement-coated antigens, these findings raise the possibility that complement-dependent antigen recognition may contribute to MZB cell activation and subsequent T cell engagement in hypertension. While the identity of the relevant antigens remains unresolved, this framework provides a mechanistic link between complement activation, B cell antigen recognition, and T cell-dependent hypertensive pathology.

Abedini et al^15^ reported that the immune microenvironment of human kidneys with CKD is enriched for B cell receptor signalling and antigen processing and presentation. Our findings extend these observations by identifying MZB-like cells as the specific subset exhibiting antigen-dependent activation in the context of hypertension. We demonstrate increased Nur77 expression, reduced BCR diversity, and clonal expansion of MZB cells in the spleens of hypertensive mice, where over half of all clones from hypertensive mice expressed heavy chain variable regions from the IGHV1 family. Thus, pro-hypertensive antigens may be selectively recognised by IGHV1-enriched BCRs on MZB cells.

Such antigen-specific activation and clonal expansion supports a mechanistic link between hypertension and autoreactive B cell responses. Together with the increase in antigen- presentation observed here and in previous studies^8, 13, 15^, these findings suggest that targeted immunomodulatory approaches, such as antigen-specific tolerization, may represent a feasible therapeutic strategy for hypertension treatment, contingent of the identification of the relevant (auto)antigens. Future studies could leverage advances in structural modelling to predict the structure of expanded IGHV1 BCRs, thereby facilitating identification of candidate pro-hypertensive antigens and informing the development of precision immunotherapies.

In conclusion, this study identifies MZB cells as the principal B cell subset that is selectively activated and expanded in murine and human hypertension. We provide the first evidence of hypertension-associated, clonally restricted BCR repertoires within this compartment, notably enriched for IGHV1 family sequences suggesting antigen-dependent activation. Our findings further implicate MZB cells in antigen-driven immune responses that amplify pathogenic T cell signalling. Importantly, these insights highlight a path toward highly targeted immunomodulatory strategies, including antigen-specific tolerization. Collectively, our findings establish MZB cells as key immunological drivers of hypertension and provide a foundation for precision immune-based approaches aimed at restoring cardiovascular homeostasis.

## 6. Author contributions

Conceptualization: H.F.G., M.J., G.R.D., A.V.; Methodology (flow cytometry and multiomic sequencing): H.F.G., M.J., M.L., A.V., A.H., M.G.L.; Methodology (tissue preparation): H.F.G., M.J., M.L., V.T., B.W., J.N.R., T.A.G.H., H.D., A.V.; Methodology (surgeries and blood pressures): H.F.G., M.L., H.D.; Formal analysis: H.F.G. and A.V.; Writing (original draft): H.F.G.; Writing (review and editing): M.G.L, M.J., A.B., C.G.S., T.J.G., G.R.D., A.V.

## 7. Acknowledgements

We acknowledge: the Bioimaging Platform, La Trobe University for flow cytometry; the Genomics Platform, La Trobe University for single-cell sequencing services; and the La Trobe Animal Research and Teaching Facility staff for their assistance with animal husbandry. Schematic figures were created with BioRender.com.

## 9. Code availability

The code used in this study is publicly available through the individual package documentation files.

## 10. Data availability

Raw and processed data generated here from the single-cell RNA sequencing and single-cell VDJ sequencing have been deposited in Gene Expression Omnibus (accession number will be made available following peer-review acceptance). Sequencing data from healthy, diabetic and hypertensive human kidneys are available for download at GSE211785^15^.

