## Supplementary material for "Antigen-dependent activation of marginal zone B cells amplifies hypertension"

### 1. Expanded Materials and Methods

#### Induction of hypertension

Mice were randomly assigned to hypertensive or normotensive groups. Hypertension was induced via subcutaneous infusion of angiotensin II (0.7 mg/kg/day) using osmotic minipumps (Alzet Model 2004, USA). Control mice received vehicle (0.1% acetic acid in saline). Infusions were maintained for either 14 or 28 days, as previously described[1]. Following pump implantation, mice were allowed to recover in warmed, clean IVC cages containing only littermates of the same sex and treatment group.

#### Systolic blood pressure recordings

Systolic blood pressure (SBP) was measured using tail-cuff plethysmography (MC4000 Multichannel System; Hatteras Instruments, USA), as previously described[2]. Briefly, mice were acclimated to the procedure on two separate occasions prior to baseline recordings. SBP was recorded weekly for 14 or 28 days following osmotic minipump implantation. Only clear, artefact-free traces were included in analyses. To minimize stress-induced variability, male and female mice were recorded separately on the same day.

#### Tissue collection and sample preparation

Bone marrow, spleen, blood and kidney samples were harvested from vehicle- and angiotensin II-infused mice for preparation into single-cell solutions. Mice were euthanised by carbon dioxide asphyxiation, followed immediately by cardiac puncture to collect blood samples after administration of enoxaparin sodium (100 µL; 400 IU; Clexane®, Auckland, NZ). Mice were then perfused with phosphate buffered saline (PBS), and tissues were collected in ice cold sterile PBS.

Bone marrow was flushed from femoral and tibial bones and spleens were mechanically dissociated. These were then passed through 70 µm sterile cell strainers and treated with red blood cell (RBC) lysis buffer (0.15 M $NH_4Cl$, 0.01 M $KHCO_3$, 6.0 mM EDTA, $dH_2O$) for 5 min at room temperature. Blood samples were centrifuged at 2,000x *g* for 10 min at 4°C. The plasma was removed, and the cell pellet was resuspended in RBC lysis buffer for 5 min, twice. Live cell counts were obtained using trypan blue exclusion and a Countess™ Automated Cell Counter (Invitrogen). Final suspensions were adjusted to $10^7$ live cells/mL in PBS and kept on ice until downstream processing.

Kidneys were halved, minced with scissors and digested in PBS containing collagenase type XI (0.16 mg/mL), collagenase type I-S (0.18 mg/mL) and hyaluronidase (0.03 mg/mL; Sigma-Aldrich, USA) for an hour at 37°C. Digested tissue was passed through a 70 μm filter, centrifuged at 450x *g* for 10 min at 4°C and resuspended in 40% isotonic Percoll™ solution (GE Healthcare Life Science, UK). A 60% Percoll™ underlay was added, and samples were centrifuged at 1385x *g* for 25 min at 25°C (no brakes). Mononuclear cells were collected from the interface, washed with PBS, and resuspended for flow cytometry.

**High-dimensional flow cytometry**

Single-cell suspensions were loaded onto 96-well microplates. Live cells were stained with LIVE/DEAD™ Fixable Blue Dead Cell Stain (1:1000 dilution in PBS; ThermoFisher Scientific) for 15 min at 4°C. Samples were washed with FACS buffer (0.5% bovine serum albumin (BSA; Sigma-Aldrich, USA) in PBS), spun and resuspended in a cocktail of fluorescently labelled antibodies against cell surface markers for 25 min at 4°C (**Supplementary Table 1)**. Single-stain controls were prepared from spleen and bone marrow aliquots.

Cells were fixed and permeabilized using eBioscience™ Fixation/Permeabilization Diluent (Invitrogen) for 30 minutes at 4°C, then stained with intracellular antibodies (**Supplementary Table 1**) for 15 min at room temperature. After final washes, cells were resuspended in 1% formalin in FACS buffer and stored at 4°C until analysis.

Flow cytometry was performed using a BD FACSymphony™ A3 analyser (BD Bioscience, San Jose, CA, USA) using the FACSDiva software v9. Data quality was ensured by removing events with unstable acquisition or flow rate using FlowAI[3] in FlowJo v10.10.0. Cleaned FCS files containing only live, non-myeloid cells were batch-corrected using the cyCombine workflow[4] with logicle transformation[5]. Batch corrected files were gated in FlowJo v10.10.0 and population counts were exported for statistical analyses. Gating strategies and population summaries are provided in **Supplementary Table 2**.

**Single-cell multiomic (RNA+VDJ) sequencing of spleen and bone marrow**

Bone marrow and spleen samples from 28 day-infused mice (n=12 per tissue) were harvested for single-cell RNA sequencing (scRNAseq) and single-cell V(D)J sequencing (scVDJseq). Each group included 3 male and 3 female mice infused with either vehicle or angiotensin II (**Supplementary Fig. 1**). Samples were dissociated into single-cell suspensions under DNAse- and RNAse-free conditions and loaded onto 96-well microplates, including additional wells for unstained and compensation controls.

Cells were incubated with Fc block (1:500 dilution in FACS buffer, BioLegend, USA) for 5 min at room temperature. Samples were indexed using TotalSeq™-C cell surface hashtags (BioLegend, USA) (**Supplementary Table 3**) diluted to 1:2000 in FACS buffer and centrifuged at 14,000 × g for 10 min at 4°C to remove antibody conjugates. Samples were then stained with an antibody master mix for fluorescence-activated cell sorting (FACS), including markers for nucleated, metabolically active and live cells (**Supplementary Table 4**), then filtered through 40 µm sterile cell strainers. Spleen samples were additionally stained with anti-CD19 and anti-CD23 antibodies to enrich for B-1 cells ($CD19^{+}CD23^{-}$), which are a minor splenic B cell population[6]. For spleens, viable B-1 cells were sorted into one tube, and all other viable cells were sorted into a second tube, prior to recombination to double the B-1 cell proportion. For bone marrow, all viable cells were sorted into a single tube. Sorted cells were then manually counted using trypan blue stain and a hemocytometer.

Samples from the same treatment group were pooled together at equal quantities and final manual cell counts were performed. Each pool contained 3 male and 3 female samples of the same tissue type. Pooled samples were placed on ice and immediately processed for single-cell RNA sequencing (scRNAseq) and single-cell V(D)J sequencing (scVDJseq) at the Genomics Platform, La Trobe University.

Gel-Bead-in-Emulsion (GEM) generation was performed using the Chromium Next GEM Chip G (PN-1000127) and Chromium controller (10X Genomics, VIC, AUS). Libraries were prepared using the Chromium Next GEM Single Cell 5′ Kit v2 (PN-1000265) and sequenced on a NovaSeq Illumina system (Illumina, Scoresby, VIC, AUS).

Gene expression libraries were generated using the CellRanger v7.0 multi-library pipeline (10X Genomics), mapped to the mouse mm10 reference transcriptome. This yielded 12 count matrices across bone marrow and spleen samples. FASTQ files were reconstructed from these matrices, BCR sequences were then extracted using sample-specific barcodes and mapped to the GRCm38 Mouse V(D)J Reference 7.0.0.

**Single-cell multiomic analyses**

Samples were annotated with treatment, sex and animal number metadata in R (version 4.2.1) using the Seurat package (v4.1.1). Mitochondrial gene expression and cell cycle scores were calculated using "PercentageFeatureSet" and "CellCycleScoring" functions, respectively[7-9]. Cells with <200 genes or >10% mitochondrial gene expression were excluded. Doublets were identified using DoubletFinder (v2.0.3), assuming a ~6.1% doublet rate based on ~$1.2 \times 10^4$ loaded cells per group[10] .

Bone marrow and spleen cells expressed an average of 2,151 and 1,627 genes with average gene counts of 9,086 and 4,202, respectively (**Supplementary Fig. 2**). Raw RNA counts were normalised using SCTransform (SCT), excluding mitochondrial gene expression and cell cycle phase from regression.

Principal component analysis (PCA) was performed on SCT-normalised data, and the first 30 PCA vectors were used for uniform manifold approximation and projection (UMAP) graph reductions. Cells were then clustered into populations using a standard resolution of 0.8.

To improve community detection, Leiden-based clustering was performed using Monocle3 (v1.2.9) on SCT-normalised counts[11-14]. UMAP reduction graphs were generated, and clusters defined using automatic resolution selection. Cluster robustness was validated using Seurat (v4.1.1), requiring $\geq$3 differentially expressed genes per cluster. Monocle3 cluster assignments and UMAP embeddings were then transferred to the original Seurat object for downstream analyses.

Cell-type predictions were performed using SingleR (v1.10.0)[15] and reference data from the Immunological Genome Project via the celldex (v1.6.0) package[15]. The top 200 differentially expressed genes (log fold change $\geq$ 0.25) were used for label assignment. Predicted labels were manually confirmed or revised using known transcript markers from the literature.

Pseudotime trajectory analysis was performed using Monocle3 (v1.2.9) to investigate transcriptional differentiation during B cell maturation. An unsupervised principal graph was constructed via reversed graph embedding of transcriptomic data, allowing cells to be ordered along a trajectory and assigned pseudotime values.

Spatial autocorrelation analysis was then used to identify genes that varied significantly along the pseudotime trajectory. Genes with a “Moran’s I” q-value $< 0.05$ were considered significantly variable and grouped into modules using a resolution of 0.001. Gene sets were retrieved from the Molecular Signature Database via the msigdbr package (v7.5.1)[16, 17], and

GO enrichment analysis was performed using the clusterProfiler package (v4.4.4)[18, 19], applying a similarity cutoff of 0.5 and a q-value threshold of < 0.05. Selected GO terms were visualised using ggplot2 (v3.5.1)[20]; full results are provided in **Supplementary files 3,4**.

Transcriptional data was integrated with antigen receptor sequencing (scVDJseq) to identify B cell populations with shared B cell receptor (BCR) sequences. Productive/functional BCRs from were identified using the Immcantation framework (Docker container)[21, 22]. BCR clonotypes were defined via unsupervised spectral clustering of BCR sequences based on V and J gene similarity using scoper (v1.2.1)[23, 24]. Diversity and species richness metrics were calculated and visualised using alakazam (v1.2.1)[25]. Scaled frequencies of variable (V) gene segments from immunoglobulin heavy (IGH) and immunoglobulin light kappa/lambda (IGK/IGL) chains were extracted using scRepertoire (v1.7.2)[26] and analysed via two-way ANOVA with Tukey's *post-hoc* test to detect biases in V region usage.

Inference and visualisation of intercellular communication was performed using CellChat (v2.1.2)[27] which compares the expression of ligand-receptor pairs between different groups and/or clusters. Weighted interactions were used to infer changes in communication strength using the 'triMean' average gene expression method. Dominant senders versus receivers were identified by calculating centrality scores. Manifold and classification learning were used to identify changes in overall- and cluster-specific signalling pathways.

**Single-cell and single-nuclear RNA sequencing of human kidneys**

Multiomic sequencing data of healthy, diabetic and hypertensive human kidneys were downloaded from GSE211785[28]. For continuity, UMAP coordinates and metadata cluster annotations from the original study were retained[28]. Prior to normalisation of raw RNA counts using SCTransform, samples from diabetic kidney disease patients, single-cell ATAC and spatial RNA sequencing were excluded. A total of 15 healthy (60% male, aged: 57.9 ± 13.5

years) and 11 hypertensive chronic kidney disease (80% male, aged: 69.5 ± 12.6 years) samples remained for downstream analyses. B cell labels were manually confirmed using known transcript markers from the literature and inference of intercellular communication pathways was performed as per our mice data.

### 2. Supplementary tables

#### Table S1: High-dimensional (spectral) flow cytometry antibodies

| Target (anti-mouse) | Clone | Catalogue number | Supplier | Flurophore | Final concentration (mg/mL) |
|---|---|---|---|---|---|
| B220 | RA3-6B2 | 103237 | BioLegend | BV570 | 0.0004 |
| CD23 | B3B4 | 101607 | BioLegend | PE | 0.0004 |
| CD11b | M1/70 | 101242 | BioLegend | BV711 | 0.0008 |
| CD138 | 281-2 | 740240 | BD Bioscience | BUV395 | 0.0008 |
| CD21 | 7E9 | 123421 | BioLegend | BV421 | 0.0008 |
| CD24 | 30-F1 | 752795 | BD Bioscience | BUV563 | 0.0008 |
| CD25 | PC61 | 569600 | BD Bioscience | BUV737 | 0.0008 |
| CD27 | LG.3A10 | 741518 | BD Bioscience | BUV661 | 0.0008 |
| CD3 | 500A2 | 560771 | BD Bioscience | V500 | 0.0008 |
| CD4 | GK1.5 | 612900 | BD Bioscience | BUV805 | 0.0008 |
| CD45 | 30-F11 | 103116 | BioLegend | APC-Cy7 | 0.0008 |
| CD5 | 53-7.3 | 25-0051-81 | eBioscience | PE-Cy7 | 0.0008 |
| CD69 | H1.2F3 | 104541 | BioLegend | BV650 | 0.0008 |
| CD8a | 53-6.7 | 100734 | BioLegend | PerCP-Cy5.5 | 0.0008 |
| CD9 | KMC8 | 740886 | BD Bioscience | BV786 | 0.0008 |
| Foxp3 (intracellular) | FJK-16s | 35-5773-82 | eBioscience | PE-Cy5.5 | 0.0008 |
| Nur77 (intracellular) | 12.14 | 566735 | BD Bioscience | AF647 | 0.0008 |
| Sca-1 | E13-161.7 | 751352 | BD Bioscience | BUV615 | 0.0008 |
| TCR beta chain | H57-597 | 747006 | BD Bioscience | BV750 | 0.0008 |
| TCR gamma-delta | GL3 | 750412 | BD Bioscience | BUV496 | 0.0008 |
| CD19 | 6D5 | 115528 | BioLegend | AF700 | 0.002 |
| CD34 | MEC14.7 | 119329 | BioLegend | PE-Dazzle | 0.002 |
| IgD | 11-26c(11-26) | 11-5993-81 | eBioscience | FITC | 0.002 |
| IgM | RMM-1 | 406523 | BioLegend | BV605 | 0.002 |
| Viability stain (LD fixable blue) | N/A | L23105 | ThermoFisher | BUV450 | N/A |

#### Table S2: Identifiable populations from spectral flow cytometry panel

| Target (anti-mouse) | Clone | Catalogue number | Supplier | Flurophore | Final concentration (mg/mL) |
|---|---|---|---|---|---|
| B220 | RA3-6B2 | 103237 | BioLegend | BV570 | 0.0004 |
| CD23 | B3B4 | 101607 | BioLegend | PE | 0.0004 |
| CD11b | M1/70 | 101242 | BioLegend | BV711 | 0.0008 |
| CD138 | 281-2 | 740240 | BD Bioscience | BUV395 | 0.0008 |

| | | | | | |
|---|---|---|---|---|---|
| CD21 | 7E9 | 123421 | BioLegend | BV421 | 0.0008 |
| CD24 | 30-F1 | 752795 | BD Bioscience | BUV563 | 0.0008 |
| CD25 | PC61 | 569600 | BD Bioscience | BUV737 | 0.0008 |
| CD27 | LG.3A10 | 741518 | BD Bioscience | BUV661 | 0.0008 |
| CD3 | 500A2 | 560771 | BD Bioscience | V500 | 0.0008 |
| CD4 | GK1.5 | 612900 | BD Bioscience | BUV805 | 0.0008 |
| CD45 | 30-F11 | 103116 | BioLegend | APC-Cy7 | 0.0008 |
| CD5 | 53-7.3 | 25-0051-81 | eBioscience | PE-Cy7 | 0.0008 |
| CD69 | H1.2F3 | 104541 | BioLegend | BV650 | 0.0008 |
| CD8a | 53-6.7 | 100734 | BioLegend | PerCP-Cy5.5 | 0.0008 |
| CD9 | KMC8 | 740886 | BD Bioscience | BV786 | 0.0008 |
| Foxp3 (intracellular) | FJK-16s | 35-5773-82 | eBioscience | PE-Cy5.5 | 0.0008 |
| Nur77 (intracellular) | 12.14 | 566735 | BD Bioscience | AF647 | 0.0008 |
| Sca-1 | E13-161.7 | 751352 | BD Bioscience | BUV615 | 0.0008 |
| TCR beta chain | H57-597 | 747006 | BD Bioscience | BV750 | 0.0008 |
| TCR gamma-delta | GL3 | 750412 | BD Bioscience | BUV496 | 0.0008 |
| CD19 | 6D5 | 115528 | BioLegend | AF700 | 0.002 |
| CD34 | MEC14.7 | 119329 | BioLegend | PE-Dazzle | 0.002 |
| IgD | 11-26c(11-26) | 11-5993-81 | eBioscience | FITC | 0.002 |
| IgM | RMM-1 | 406523 | BioLegend | BV605 | 0.002 |
| Viability stain (LD fixable blue) | N/A | L23105 | ThermoFisher | BUV450 | N/A |

### Table S3: Hashtags used to index samples for single-cell sequencing

| Product | Catalogue number | Barcode sequence |
|---|---|---|
| TotalSeq™-C0301 anti-mouse Hashtag 1 Antibody | 155861 | ACCCACCAGTAAGAC |
| TotalSeq™-C0302 anti-mouse Hashtag 2 Antibody | 155863 | GGTCGAGAGCATTCA |
| TotalSeq™-C0303 anti-mouse Hashtag 3 Antibody | 155865 | CTTGCCGCATGTCAT |
| TotalSeq™-C0304 anti-mouse Hashtag 4 Antibody | 155867 | AAAGCATTCTTCACG |
| TotalSeq™-C0305 anti-mouse Hashtag 5 Antibody | 155869 | CTTTGTCTTTGTGAG |
| TotalSeq™-C0306 anti-mouse Hashtag 6 Antibody | 155871 | TATGCTGCCACGGTA |

### Table S4: Antibodies and dyes used to sort cells for single-cell sequencing

| Target (anti-mouse) | Clone | Catalogue number | Supplier | Flurophore | Final concentration |
|---|---|---|---|---|---|
| CD16/32 (Fc block) | 93 | 14-0161-81A | eBioscience | Unconjugated | 0.001 (mg/mL) |
| Vybrant™ DyeCycle™ Ruby (nucleated dye) | NA | V10273 | Invitrogen | NA | 5 (μM) |
| Calcein AM Blue (metabolic viability dye) | NA | 65-0855-39 | Invitrogen | NA | 0.1 (μM) |
| SYTOX™ Green (dead cell dye) | NA | S34860 | Invitrogen | NA | 0.03 (μM) |
| CD19 | 6D5 | 115520 | BioLegend | PE-Cy7 | 0.0004 (mg/mL) |
| CD23 | B3B4 | 101607 | BioLegend | PE | 0.0004 (mg/mL) |

**Table S5: Gene markers and references for overall cluster identification**

| Population | Gene marker(s) | References |
|---|---|---|
| Pro-neutrophils | *Elane, Cebpe* | 29 |
| Neutrophils | *Retnlg, Mmp9, Wfdc21* | 29 |
| Erythroid primed monocyte-erythrocyte progenitors (E-MEPs) | *Tfrc* (CD71), *Gypa* (CD235a), *Klf1, Tmod1, Ank1* | 30 |
| Granulocyte-monocyte progenitors (GMPs) | *Cd34, Kit, Angpt1* | 31 |
| Pro-monocytes | *Plppr3, Flt3, Csf1r* (CD115), *Ly6c2* | 32 |
| Monocytes | *Ms4a6c, Fn1* | 33 |
| Basophils | *Cd200r3, Gata2, Fcer1a* | 34 |
| Dendritic cells (DCs) | *Siglech, Ccnd1, Cox6a2* | 35 |
| Natural killer cells (NKs) | *Ncr1, Gzma, Klra8* | 36 |
| T cells | *Cd3e, Cd3g, Themis* | 37 |
| B cells | *Cd79a, Cd19, Ms4a1* | 38-41 |

**Table S6: Gene markers and references for B cell subpopulation identification**

| Population | Gene marker(s) | References |
|---|---|---|
| Pre-pro B | *Ly6d (low)* | 42 |
| Pro-B | *Ly6d (low), Rag1, Rag2, Vpreb1, Vpreb2, Vpreb3, Il7r* | 42-46 |
| Pre-B cell receptor dependent (Pre-BCRd) | *Ly6d (low), Vpreb1, Vpreb3, Mki67, Pclaf ,Top2a, Il7r* | 38, 42, 43, 46-48 |
| Pre-B cell receptor independent 1 (Pre-BCRi 1) | *Iglc1, Iglc2, Iglc3, Il7r* | 38, 46, 49-51 |
| Pre-B cell receptor independent 2 (Pre-BCRi 2) | *Iglc1, Iglc2, Iglc3* | 38, 46, 49-51 |
| Transitional B cells 1 (Tr1) | *Iglc1, Iglc2, Iglc3, Tnfrsf13c, Cd24a, Ms4a1* | 38, 52-55 |
| Transitional B cells 2 (Tr2) | *Iglc1, Iglc2, Iglc3, Tnfrsf13c, Cd24a, Ms4a1* | 38, 52-55 |
| Circulating B-2 cells (Circ. B-2) | *Iglc1, Iglc2, Iglc3, Fcer2a* (CD23), *Cr2* (CD21) | 56, 57 |
| Follicular B-2 cells (FO B-2) | *Iglc1, Iglc2, Iglc3, Fcer2a* (CD23), *Cr2* (CD21), *Klf2* | 56, 57 |
| Marginal zone B cells (MZB) | *Iglc1, Iglc2, Iglc3, Fcer2a* (low) (CD23), *Cr2* (CD21), *Mzb1* | 56, 57 |
| B-1 cells | *Iglc1, Iglc2, Iglc3, Mzb1* | 56, 57 |
| Germinal centre cells (GC) | *Iglc1, Iglc2, Iglc3, Aicda* | 58 |

| | | |
|---|---|---|
| Memory B cells | *Iglc1, Iglc2, Iglc3, Bach2, Dock2* | 59, 60 |
| Plasma cells | *Tnfrsf17, Prdm1, Sdc1* | 38, 61, 62 |
| Anti-viral B cells | *Ifit3, Isg15, Irf7* | 63-68 |

### 3. Supplementary Figures

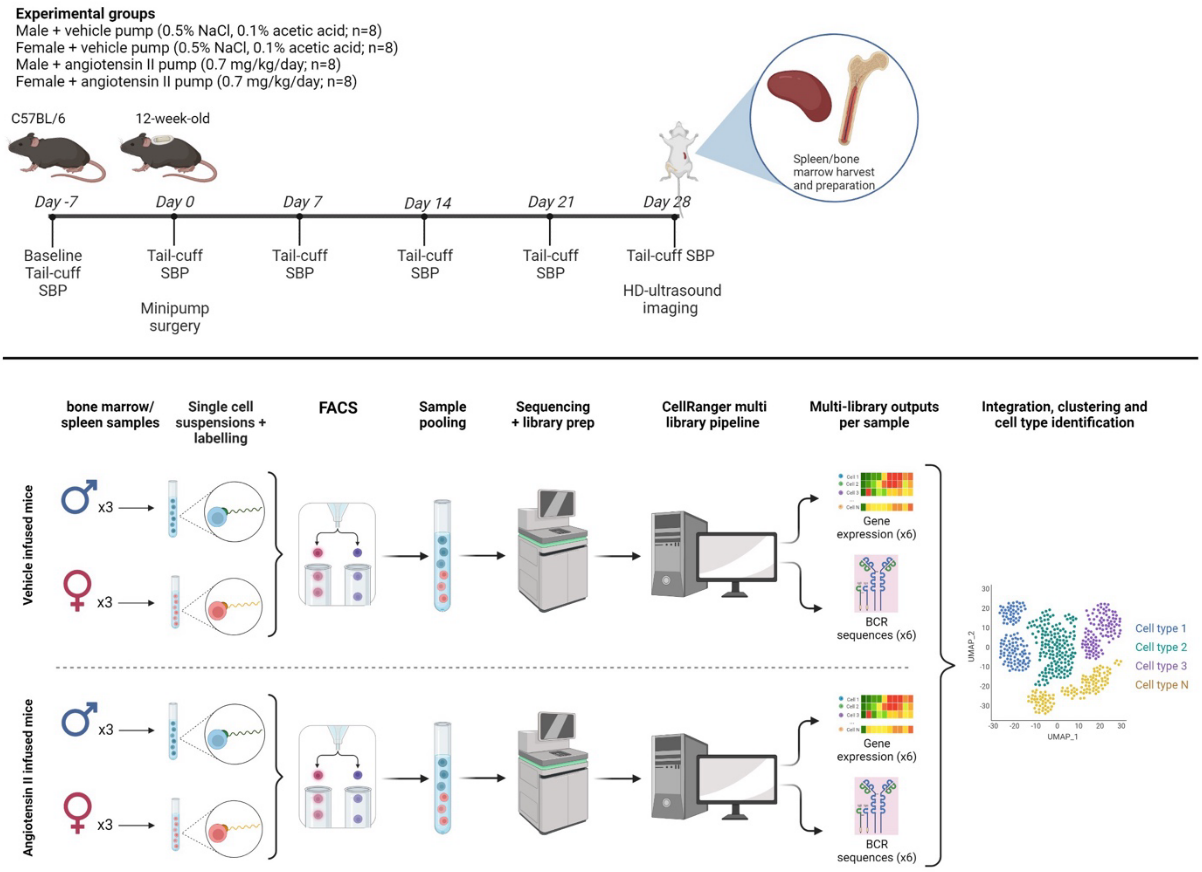


**Figure S1: Study design and multiomic sequencing pipeline.** Image created with BioRender.com

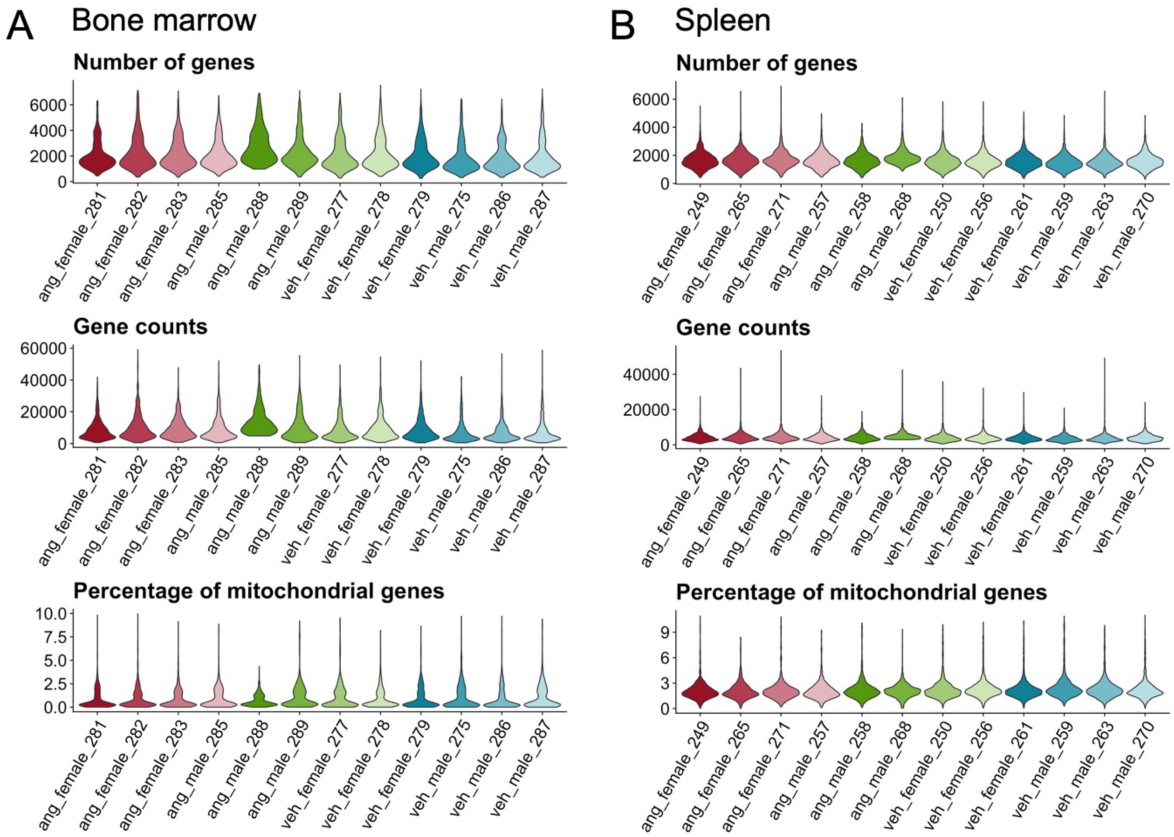


C

| Bone marrow populations | Number of cells | |
|---|---|---|
| | VEH | ANG |
| B cells | 898 | 753 |
| Basophils | 38 | 49 |
| Dendritic cells | 112 | 151 |
| Erythroid primed MEPs | 21 | 26 |
| Granulocyte-Monocyte progenitors | 65 | 79 |
| Monocyte progenitor | 113 | 107 |
| Monocytes | 647 | 495 |
| Natural killer cells | 33 | 30 |
| Neutrophil progenitors | 147 | 148 |
| Neutrophils | 3394 | 3129 |
| T cells | 83 | 150 |
| **Total** | **5551** | **5117** |

D

| Spleen populations | Number of cells | |
|---|---|---|
| | VEH | ANG |
| B cells | 3699 | 2877 |
| Basophils | 28 | 20 |
| Dendritic cells | 106 | 83 |
| Monocytes | 93 | 136 |
| Natural killer cells | 416 | 332 |
| Neutrophils | 26 | 48 |
| Stem cells | 9 | 16 |
| T cells | 3989 | 3485 |
| **Total** | **8366** | **6997** |

**Figure S2: Sequencing metrics.** Violin plots showing the total number of sequenced genes, gene counts and percentage of mitochondrial gene expression per animal from bone marrow (A) and spleen (B) samples. Tables C and D summarise the number of cells sequenced in the bone marrow and spleen, respectively, by treatment group.

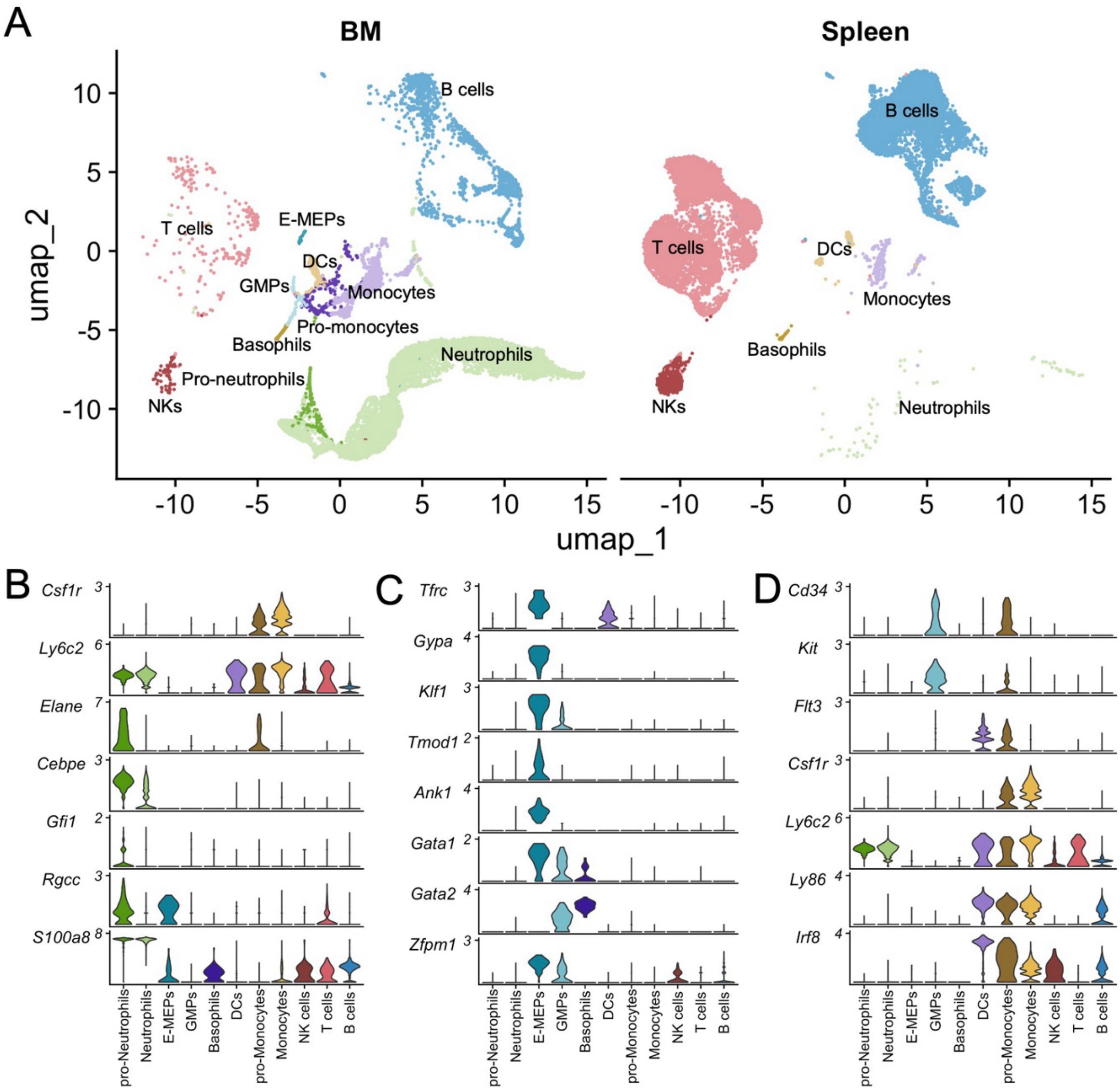


**Figure S3: Gene markers of hematopoietic cells in the bone marrow.** A) UMAP dimensional reduction showing cells sequenced in the bone marrow and spleen. Violin plots showing the expression of cell type specific genes for B) pro-neutrophils, C) E-MEPs and D) GMP/monocyte markers. BM = bone marrow; E-MEPs= erythroid primed monocyte-erythrocyte progenitors; GMPs = granulocyte-monocyte progenitors; DCs = dendritic cells; NKs = natural killer cells

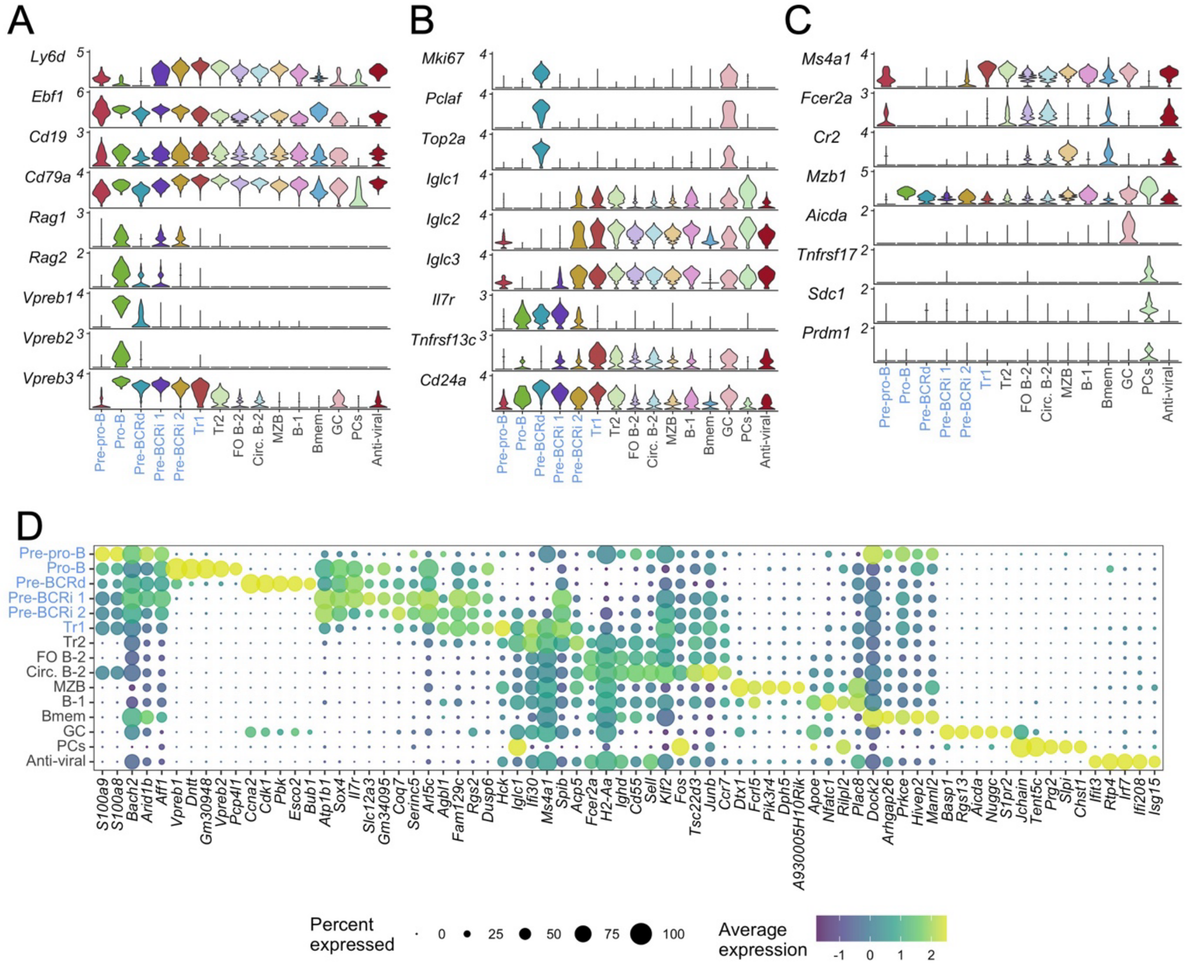


**Figure S4: Earliest committed progenitor B cells through to terminally differentiated plasma cells were captured in the spleen and bone marrow by single cell transcriptomics.** Early-stage B cell markers (A), proliferative and transitional B cell markers (B) and mature B cell markers (C) shown in violin plots. D) Top 5 uniquely expressed genes for each population where the size and colour of each dot represents the percentage of cells expressing a given gene in that cluster and its normalized expression count, respectively. Immature B cell populations found exclusively in the bone marrow are shown in cornflower blue. Pre-BCRd = pre-B cell receptor dependent cells; Pre-BCRi = pre-B cell receptor independent cells; Tr1 = transitional B cells found in bone marrow; Tr2 = transitional B cells found in spleen; MZB = marginal zone B cells; Bmem = memory B cells; GC = germinal center B cells; PCs = plasma cells; Circ. B-2 = circulating B-2 cells.

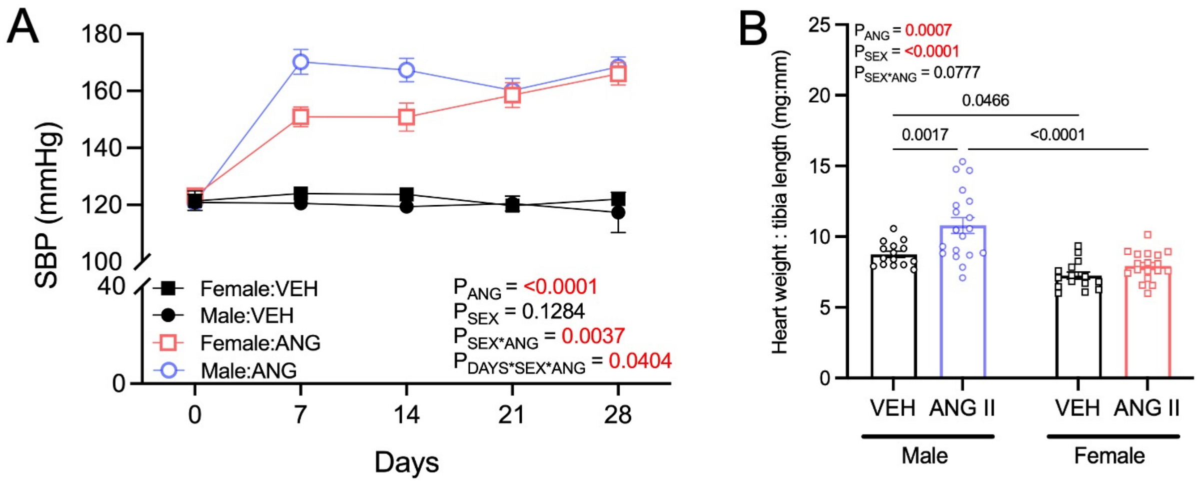


**Figure S5: Female mice have a slower response to angiotensin II and lower cardiac hypertrophy.** A) Systolic blood pressures (a) of vehicle (0.5% NaCl, 0.1% acetic acid) and angiotensin II (0.70 mg/kg/day)-infused male and female mice. B) Heart-weight to tibia length ratios of male and female normotensive and hypertensive mice.

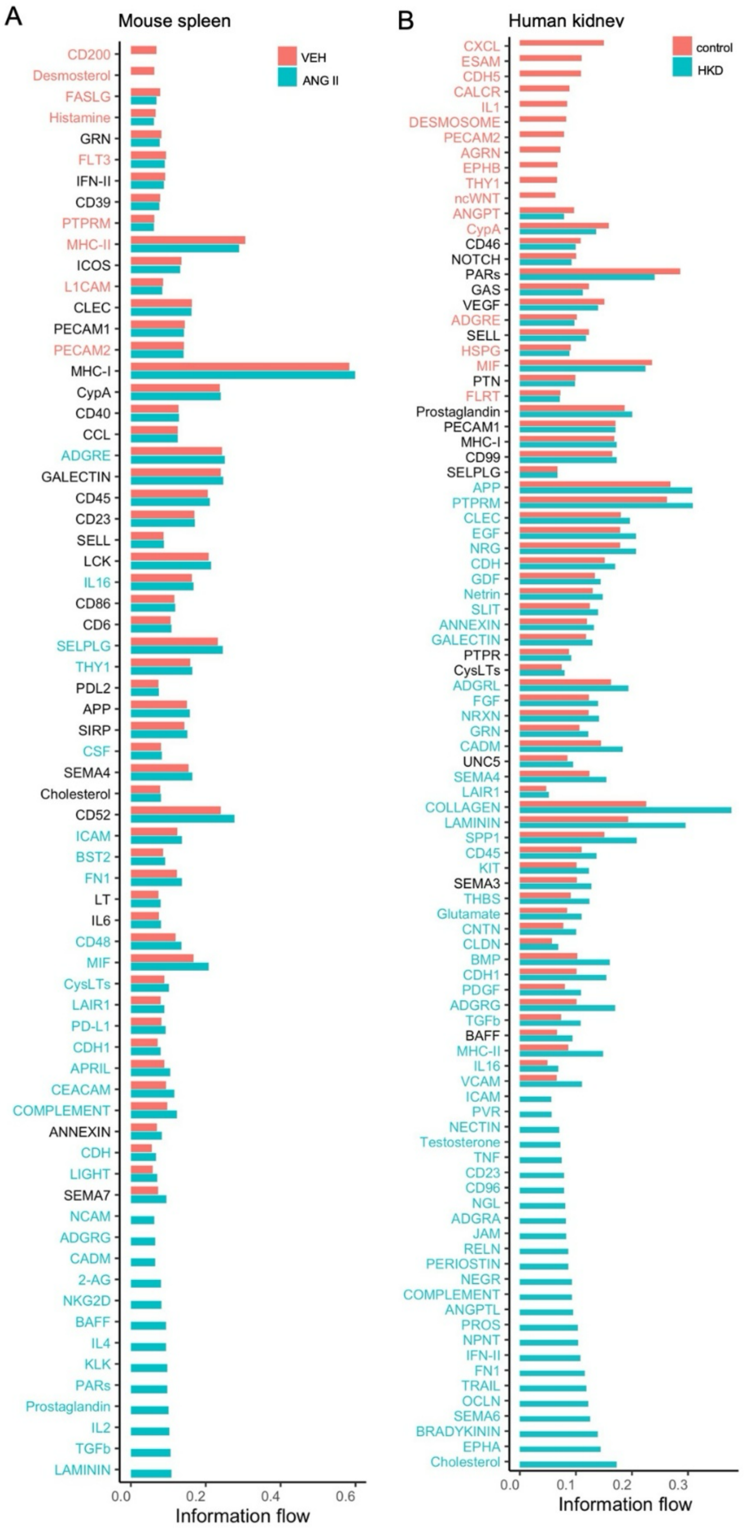


**Figure S6: Mice and humans share 11 signalling pathways upregulated during hypertension.** Comparison of specific signalling pathways enriched either in normotensive and hypertensive mice (A) and humans (B). Pathways shown in red are significantly enriched in vehicle-infused mice or control samples from normotensive humans. Pathways in blue are significantly enriched in angiotensin II-infused mice or samples from participants with hypertensive chronic kidney disease.
